# Programmable *in vivo* CRISPR activation elucidates the oncogenic and immunosuppressive role of MYC in lung adenocarcinoma

**DOI:** 10.1101/2021.12.09.471859

**Authors:** Fredrik I. Thege, Dhwani N. Rupani, Bhargavi B. Barathi, Sara L. Manning, Anirban Maitra, Andrew D. Rhim, Sonja M. Wörmann

## Abstract

Conventional genetically engineered mouse models (GEMMs) are time consuming, laborious and offer limited spatio-temporal control. Here, we describe the development of a streamlined platform for *in vivo* gene activation using CRISPR activation (CRISPRa) technology. Unlike conventional GEMMs, our model system allows for flexible, sustained and timed activation of one or more target genes using single or pooled lentiviral guides. Using *Myc* and *Yap1* as model oncogenes, we demonstrate gene activation in primary pancreatic organoid cultures *in vitro* and enhanced tumorigenic potential in *Myc*-activated organoids when transplanted orthotopically. By implementing our model as an autochthonous lung cancer model, we show that transduction-mediated *Myc* activation leads to accelerated tumor progression and significantly reduced overall survival relative to non-targeted tumor controls. Furthermore, we found that *Myc*-activation led to the acquisition of an immune suppressive “cold” tumor microenvironment. Through cross-species validation of our results using publicly available RNA/DNA-seq data sets, we were able to link MYC to a previously described, immunosuppressive transcriptomic subtype in patient tumors, thus identifying a patient cohort that may benefit from combined MYC/immune-targeted therapies. Overall, our work demonstrates how CRISPRa can be used for rapid functional validation of putative oncogenes and may allow for the identification and evaluation of potential metastatic and oncogenic drivers through competitive screening.

## Introduction

Improved technologies for introducing controlled perturbations to gene expression *in vivo* hold great promise for the interrogation of gene function and modeling of human disease. While conventional genetically engineered mouse models (GEMMs) require that a custom model be created for each individual gene-of-interest, at significant time and monetary costs, a programmable mouse model for gene activation, where one or more genes can be activated in an *ad hoc* manner, would constitute a powerful tool for cost and time efficient modeling of genetic drivers in human disease, including cancer.

In GEMMs where tumor formation is driven through tissue-specific Cre-Lox recombination, oncogenic transformation occurs concurrently in all Cre expressing cells. This is in stark contrast to how tumors form in patients, where most tumors arise from stochastic events in a small population of cells. Transduction/transfection-based tumor initiation more closely recapitulates this process as only a fraction of cells are transduced, and thus have the potential for tumor initiation.

Since its description as a programmable system for gene editing(1–3), CRISPR/Cas9 has become a ubiquitous scientific tool for gene knock out (CRISPR KO), interference (CRISPRi), and activation (CRISPRa). While systems for *in vivo* CRISPR KO have been implemented using mice conditionally expressing Cas9 (LSL-Cas9 mice)(4) and RNA guides delivered through viral transduction(5) or plasmid electroporation(6,7), the incorporation of CRISPRa in *in vivo* platforms has to date been limited. The CRISPR/Cas9 Synergistic Activation Mediator (SAM) system is a robust three-component system for CRISPRa, consisting of dCas9 fused to four tandem repeats of the herpes simplex transactivator VP16 (dCas9VP64), a transactivating fusion protein (MS2-p65-HSF1) and MS2 aptamer-loop modified single guide RNA (sgRNA(MS2))(8,9). The SAM system has been used to drive expression of individual or multiple genes using single or pooled RNA guides across a wide range of cell types *in vitro(8–11)*, and was recently implemented *in vivo(12)*.

In contrast to CRISPR KO, CRISPRa relies on stable integration of the RNA guide into the host genome for retained guide expression, sustained gene activation and tracing of guide abundance over long periods of time, e.g. during tumor formation. Therefore, for oncogene studies, lentiviral delivery(13,14) of sgRNA sequences is favorable over transient adeno-associated virus (AAV) or adenovirus systems(15–17).

Unlike standard transgene-based over-expression technologies, CRISPRa relies on recruitment of transactivators to the endogenous gene locus, and thus more faithfully retains important aspects of transcriptional diversity and regulation, limits the level of over-expression, enables isotype-specific activation, and allows for protein-coding genes that are too large for transduction/transfection and non-coding transcripts (e.g. lincRNAs) to be activated(8).

In addition to *in vivo* gene activation, a CRISPRa enabled mouse model would serve as an indefinite and versatile source of CRISPRa competent primary cells suitable for *ex vivo* expansion, screening and/or transplantation. *In vivo* CRISPRa oncogene activation elegantly takes advantage of the fact that a gene activation that leads to increased tumorigenicity or proliferative ability will be positively selected for and remain in the population. This creates the opportunity to use a CRISPRa mouse, in conjunction with prerequisite oncogenic drivers (e.g. Kras mutation, P53 loss), as a platform of *in vivo* oncogene evaluation and screening.

With this in mind, we created a model for oncogene activation consisting of a GEMM harboring conditional expression of the components necessary for CRISPRa/SAM, in combination with Cre/sgRNA-encoding lentivirus that can be introduced in a targeted manner through direct and local administration. While we hypothesize that a wide range of human tumor entities can be modeled using this approach, we describe a first proof-of-principle implementation of our system by targeting MYC and YAP1, two important oncogenes across human cancer, in pancreatic organoid cultures and in an autochthonous lung cancer model.

Using our LSL-SAM mouse model, we show successful *Myc* and *Yap1* gene activation in primary pancreas organoid cultures *in vitro*, and enhanced tumorigenicity following orthotopic transplantation of *Myc*-activated organoids. By incorporating conditional P53 loss and oncogenic Kras in our LSL-SAM platform to create an autochthonous lung cancer model, we were able to trigger gene activation and tumor formation *in vivo* using nasal instillation of Cre/sgRNA lentivirus.

Moreover, *Myc*-activated mice showed rapid onset disease progression and significantly reduced survival relative to controls. Transcriptomic analyses were consistent with MYC and YAP1-associated transcriptomic reprogramming in gene activated tumors. Consistent with work of others(18–21), we found a pronounced cold immune microenvironment in *Myc*-activated tumors. We furthermore identified *Myc* as a driver of the immune suppressive lung adenocarcinoma subtype 2 (LuAd2) in mouse and human lung tumors.

Using our *in vivo* CRISPRa/SAM platform we were able to recapitulate the tumorigenic and immune evasive role of MYC in human lung adenocarcinoma. In addition, our work provides cross species evidence that MYC is a driver of the LuAd2 subtype(22), highlighting a previously unexplored opportunity for targeted therapy in this recalcitrant subtype.

## Results

### Development of a programmable mouse model for *in vivo* gene activation

To generate a mouse model for programmable and traceable gene activation *in vivo*, we set out to create a mouse harboring conditional expression of the main components of the CRISPRa/SAM system and a lineage tracer. With this mouse we reasoned that we would be able to trigger tissue-specific gene activation through targeted delivery of lentivirus co-expressing Cre recombinase and sgRNA (Figure 1a). The LSL-SAM mouse was created by knocking in conditional, CAG-driven, loxp-stop-loxp (LSL) controlled, tricistronic expression of dCas9VP64, MS2-p65-HSF1 and mCherry in the Rosa26 (R26) mouse locus (Figure 1b-c, Supplemental Figure 1a-b). To confirm functional activity of the construct, we generated pancreatic ductal organoids from R26(LSL-SAM) and YFP reporter (R26(LSL-YFP)) mice and transduced these organoids with Cre adenovirus (AdCre) *in vitro*, which resulted in LSL cassette recombination and reporter (mCherry/YFP) expression (Supplemental Figure 1c-d). To confirm conditional transgene expression *in vivo*, we generated Ptf1a(Cre/+) R26(LSL-SAM/LSL-YFP) mice, expressing the SAM construct and YFP specifically in the pancreas epithelium. We found that pancreas acinar cell clusters from these mice co-expressed mCherry and YFP (Figure 1d) and pancreas tissues expressed nuclear dCas9VP64 (Figure 1e-f). To test our ability to trigger expression of the SAM components through transduction *in vivo*, we delivered Cre lentivirus directly to the pancreas via retrograde ductal injections in R26(LSL-SAM) mice, which resulted in cells sporadically expressing nuclear dCas9VP64 (Figure 1g).

**Figure 1.**
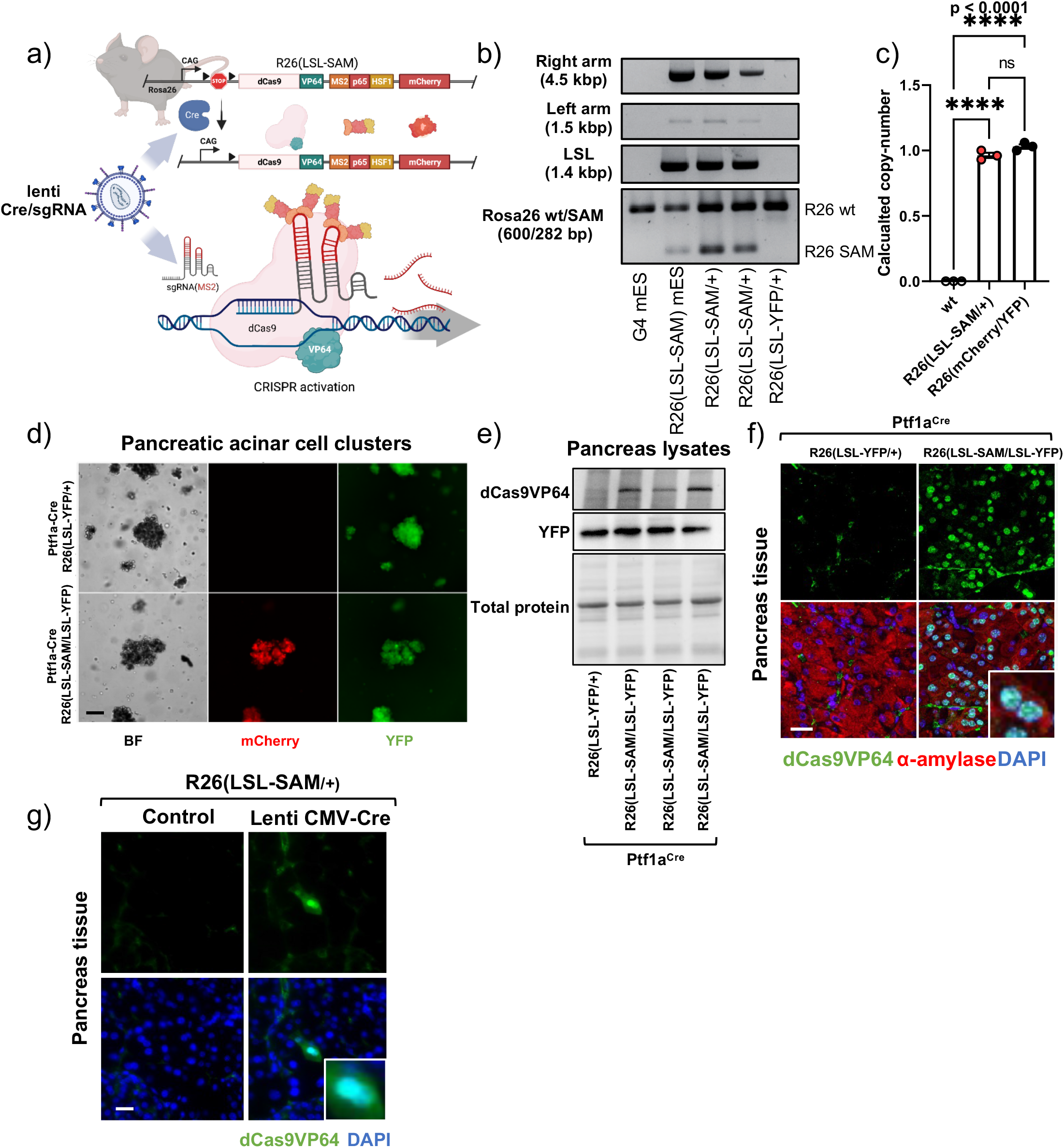
Development and validation of the LSL-SAM mouse model. **(a)** Schematic of the CRISPR activation/SAM mouse model for programmable and tissue-specific gene activation, consisting of the LSL-SAM mouse and Cre/sgRNA encoding lentivirus. **(b)** PCR confirmation of LSL-SAM Rosa26 knock-in (right and left homology arms), intact LSL cassette, and genotyping for the LSL-SAM construct in mouse embryonic stem cells and knock-in mice. **(c)** LSL-SAM copy number determination using TaqMan Copy Number assay for mCherry, calibrated to mouse DNA sample with known mCherry copy number. One-way ANOVA with Tukey’s multiple comparison test. **(d)** Fluorescent imaging of pancreatic acinar clusters isolated from Ptf1a(Cre/+) R26(LSL-SAM/LSL-YFP) mice co-expressing mCherry and YFP, indicating recombination and construct activation, scale bar 200 μm. **(e)** Immunoblot analysis showing dCas9VP64 and YFP expression in pancreatic tissues from Ptf1a(Cre/+) R26(LSL-SAM/LSL-YFP) mice and total protein as loading control, Ptf1a(Cre/+) R26(LSL-YFP/+) mouse used as control. **(f)** Immunofluorescent staining for dCas9VP64 in pancreas tissues from Ptf1a(Cre/+) R26(LSL-SAM/LSL-YFP) mice revealing nuclear dCas9VP64 localization, Ptf1a(Cre/+) R26(LSL-YFP/+) used as control, dCas9VP64 (green), α-amylase (red), DAPI (blue), scale bar 20 μm. **(g)** Sporadic nuclear expression of dCas9VP64 in pancreas tissues from R26(LSL-SAM/+) mice transduced with CMV-Cre lentivirus through retrograde ductal injection 10 days post transduction, and untreated control, dCas9VP64 (green), DAPI (blue), scale bar 20 μm.

### Functional validation of the CRISPRa/SAM mouse model in organoid transplantation models

Next, to allow for controllable induction of sgRNA and SAM construct expression *in vitro* and to test gene activation in primary cells, we designed and cloned multiple lentiviral transfer plasmids encoding for Cre or tamoxifen-inducible Cre (CreERT2), and an antibiotic selection marker (puroR, zeoR or hygroR), in addition to human U6 (hU6) promoter-driven sgRNA(MS2) (Figure 2a). We next transduced and selected pancreatic organoids established from R26(LSL-SAM) mice with CreERT2/puroR lentivirus and administered 4-hydroxytamoxifen (4OHT) to trigger Cre recombination, which resulted in SAM construct activation (Figure 2a-c). To validate programmable sgRNA-targeted gene activation, we transduced and activated LSL-SAM organoids with lentivirus encoding for sgRNAs targeting mouse *Myc* or *Yap1*, or non-targeting control (see methods for guide sequences), which resulted in 4.4 and 2.8-fold transcriptomic activation, respectively, and was confirmed with whole-mount immunofluorescence staining (Figure 2d-f). To test functional gene activation in the context of oncogenic drivers we triggered *Myc* activation in “pre-oncogenic” organoids harboring conditional loss of P53 and activating KrasG12D mutations in addition to the LSL-SAM construct: (P53(F/F) Kras(LSL-G12D/+) R26(LSL-SAM); PPKS), resulting in 3.7-fold *Myc* transcriptional activation (Figure 2g-i). *Yap1* activation yielded similar results (Supplemental Figure 2a-c). To test if *Myc* activation in the context of P53 loss and oncogenic Kras increases organoid tumorigenicity, we transplanted *Myc*-activated (PPKS/M) and non-targeted (PPKS/NT) ductal pancreatic organoids orthotopically into the pancreata of syngeneic mice (Figure 2j). We found that transplanted organoids were able to initiate tumor formation (Figure 2k) and that *Myc*-activation significantly accelerated tumor progression and death in transplanted mice (median survival 92 vs 133 days, p=0.021, Figure 2l, Supplementary Table 1).

**Figure 2.**
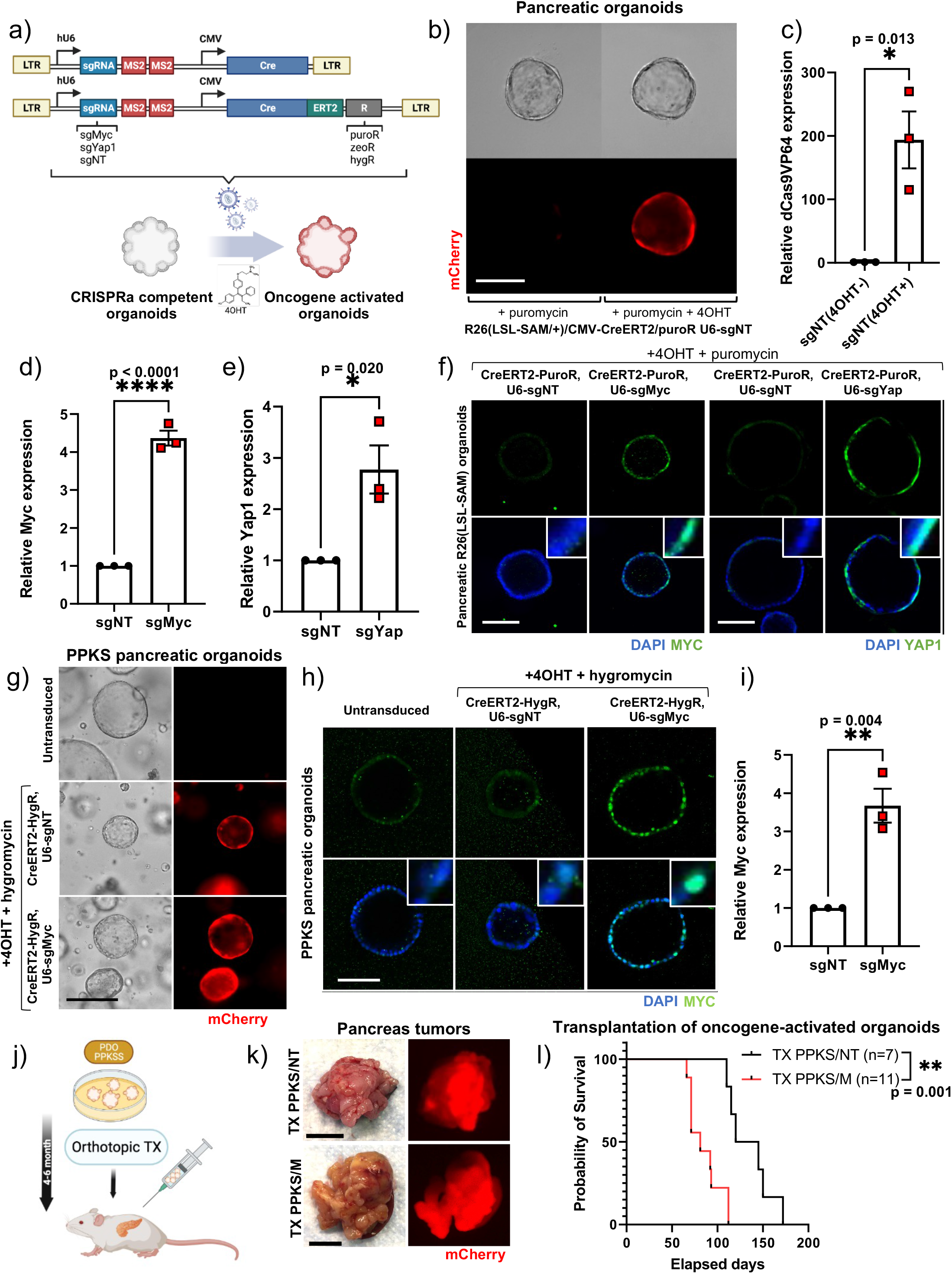
*Ex vivo* gene activation in pancreatic organoids derived from LSL-SAM/CRISPRa mice and enhanced tumorigenic potential following *Myc*-activation. **(a)** (top) Schematic of the CMV-Cre-U6-sgRNA(MS2) and CMV-CreERT2-P2A-puroR/zeoR/hygR-U6-sgRNA(MS2) lentiviral vectors, (bottom) *ex vivo* gene activation in LSL-SAM pancreatic organoids following lentiviral transduction and Cre-induction with 4-hydroxytamoxifen (4OHT). **(b)** SAM construct activation in R26(LSL-SAM) pancreatic organoids following transduction with CMV-CreERT2-P2A-puroR lentivirus, puromycin selection and Cre-induction with 4OHT. mCherry (red), scale bar 100 μm. **(c)** qPCR analysis showing expression of dCas9VP64 in pancreas organoids following transduction with CMV-CreERT2-P2A-puroR lentivirus, selection and treatment with 4OHT, relative to uninduced controls, unpaired, two-tailed, Student’s t-test. **(d)** qPCR analysis showing *Myc* gene activation in LSL-SAM pancreatic organoids transduced with *Myc*-targeted (sgMyc) and non-targeted (sgNT) CMV-CreERT2-P2A-puroR-U6-sgRNA(MS2) lentivirus. **(e)** qPCR analysis showing *Yap1* gene activation in LSL-SAM pancreatic organoids transduced with *Yap1*-targeted (sgYap) and non-targeted (sgNT) CMV-CreERT2-P2A-puroR-U6-sgRNA(MS2) lentivirus. **(f)** Wholemount immunofluorescent staining for MYC (left) and YAP1 (right) in LSL-SAM pancreatic organoids transduced with *Myc*, *Yap1* or non-targeted CMV-CreERT2-P2A-puroR-U6-sgRNA(MS2) lentivirus, MYC/YAP1 (green), DAPI (blue), scale bar 100 μm. **(g)** SAM construct activation in “pre-oncogenic” pancreatic organoids carrying conditional P53 knock-out, KrasG12D, and SAM constructs (P53(F/F), Kras(LSL-KrasG12D/+) and Rosa26(LSL-SAM); PPKS) showing mCherry expression following transduction with *Myc*-targeted or non-targeted CMV-CreERT2-P2A-hygR-U6-sgRNA(MS2) lentivirus, hygromycin selection and induction with 4OHT, and untranduced/untreated PPKS organoids as control, scale bar 100μm. **(h)** Wholemount immunofluorescent staining for MYC in PPKS pancreatic organoids transduced with *Myc*-targeted or non-targeted CMV-CreERT2-P2A-hygR-U6-sgRNA(MS2) lentivirus and untransduced controls, MYC (green), DAPI (blue), scale bar 100 μm. **(i)** qPCR analysis showing *Myc* gene activation in PPKS pancreatic organoids transduced with *Myc*-targeted CMV-CreERT2-P2A-hygR-U6-sgRNA(MS2) lentivirus relative to non-targeted control. **(j)** Schematic of orthotopic transplantation of *ex vivo* expanded and oncogene activated PPKS pancreatic organoids. **(k)** Pancreas tumors resulting from orthotopic transplantation of *Myc*-activated PPKS organoids (bottom) and non-targeted controls (top), color image (left), macroscopic mCherry fluorescence image (right), scale bar 1 cm. **(l)** Kaplan-Meier survival graph of mice orthotopically transplanted with *Myc*-activated PPKS organoids (TX PPKS/NT) and non-targeted controls (TX PPKS/NT), revealing significantly shorter survival for mice receiving *Myc*-activated organoids. Mantel-Cox (logrank) test.

Thus far, we have demonstrated the functionality of our LSL-SAM mouse and the ability to induce programmable gene activation in primary cells derived from this model. We also demonstrated the ability to generate and transplant tumorigenic organoids with programmable gene activation. This approach could straightforwardly be expanded to allow for the study of individual or pooled genetic drivers of disease using *ex vivo* expansion, manipulation, and transplantation/adoptive transfer of primary cells from the LSL-SAM mouse model.

### *In vivo* transduction-mediated tumor initiation in an autochthonous lung cancer model

To allow for programmable gene activation *in vivo*, we next developed a transduction approach using Cre/sgRNA encoding lentivirus. First, we created an assay for functional Cre-lentivirus titer quantification. To this end, we generated a color-switching Cre-reporter cell line and a protocol for functional Cre-lentivirus quantification (Figure 3a-b, see methods). To achieve sufficiently high lentiviral titers for *in vivo* use, lentiviral supernatants were concentrated using PEG/NaCl-mediated precipitation. Using this approach, we were able to achieve ultracentrifugation-level performance, at greatly increased throughput and substantially reduced cost, routinely achieving functional Cre-titers of 3e9TU/ml (Figure 3c-d). To test transduction *in vivo*, we used nasal instillation to administer lentivirus directly to the mouse lung (Figure 3e). To test whether lentiviral transduction results in significant stable genomic incorporation and transgene expression, we treated mice with CMV-eGFP encoding lentivirus. We detected eGFP-expressing cells in the lung parenchyma after 10 days and 4 weeks, indicating genomic integration and stable expression (Figure 3f). We next set out to determine the dose-response relationship of Cre lentivirus and recombination in the lung. CMV-Cre encoding lentivirus was administrated at various doses to YFP reporter mice (R26(LSL-YFP)) and the fraction of YFP-expressing cells in the lung was determined after 10 days. We found a rough dose-response relationship between viral dose and the number of YFP expressing cells in the lung parenchyma, with significant mouse-to-mouse variability (Figure 3g-h, Supplementary Table S2). We determined that a viral dose of 1e8 TU gave rise to a maximum level of transduction-mediated recombination (~80 cells/mm^2^). In addition, we compared the performance of CMV and Ef1a promoter in driving Cre-expression in the lung and found that they performed equivalently (Figure 3i, Supplementary Table S2). Taking these findings into consideration, we generated lung tumors by transducing (P53(F/F) Kras(LSL-G12D/+) R26(LSL-YFP) (PPKY) mice with CMV-Cre lentivirus through nasal instillation at a dose of 1e8TU. The resulting tumors were found to be positive for Cre recombination, and to harbor stable genomic integration of the viral payload (Figure 3j, Supplemental figure 3a-c). Cre expression was detected in normal lung tissues 10 days following CMV-Cre lentivirus transduction and in transduction-initiated tumors (Figure 3k).

**Figure 3.**
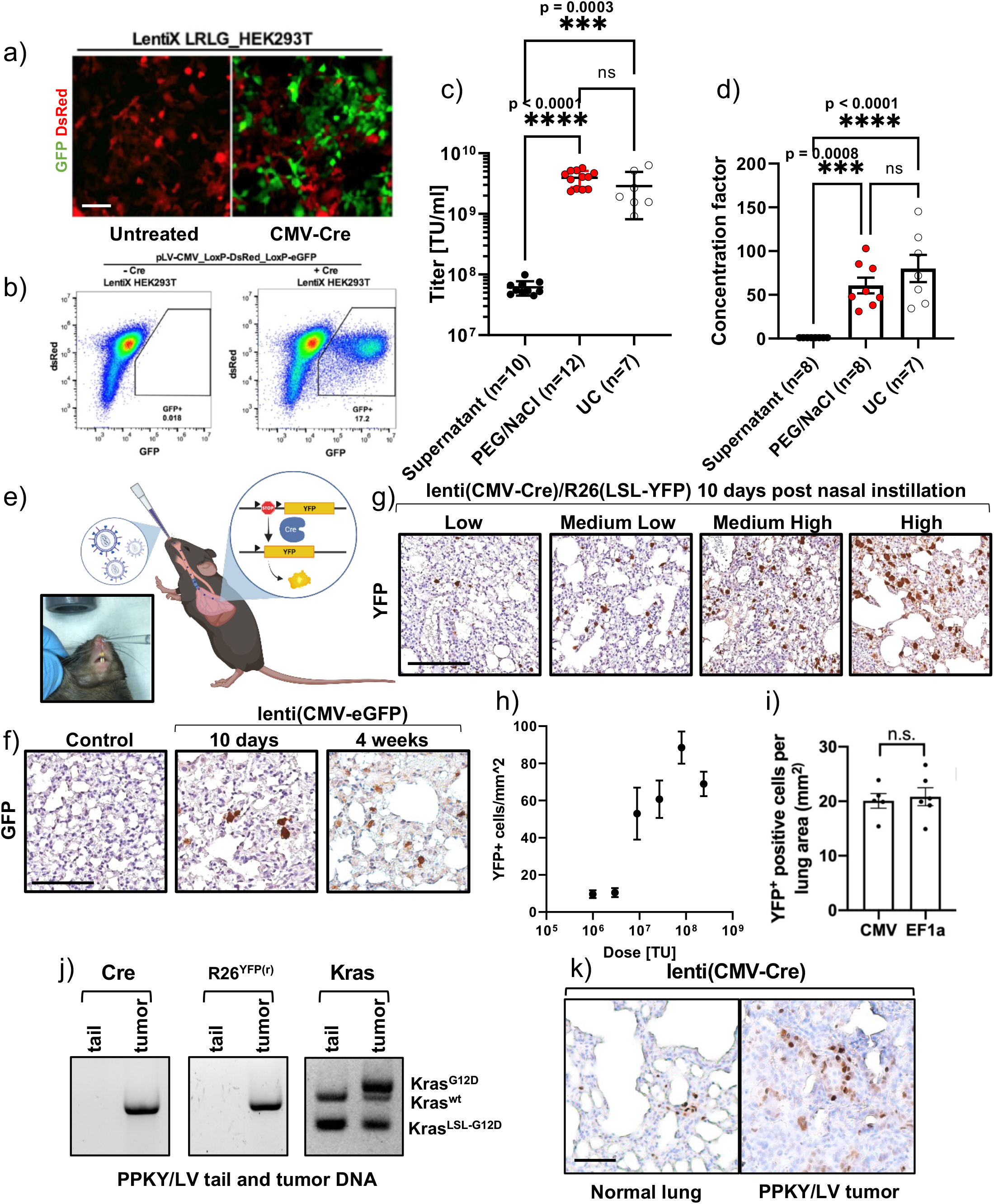
Optimization of lentiviral production, quantification and transduction for *in vivo* gene activation. **(a)** Color-switching Cre-reporter HEK293T cell line (LRLG) that switches from DsRed (left) to GFP (right) following Cre-mediated recombination, scale bar 100 μm. **(b)** Flow cytometric lenti Cre-titer quantification by determining the fraction of color-switched (GFP expressing) LRLG cells, untransduced sample (left), LRLG cells transduced with CMV-Cre lentivirus (right). **(c)** Comparison of PEG/NaCl-precipitation and ultracentrifugation (UC) for concentrating Cre lentivirus, one-way ANOVA with Tukey’s multiple comparison test. **(d)** Precipitation and ultracentrifugation concentration factor relative to viral supernatant, theoretical concentration factor 100x (initial volume/final volume = 100), one-way ANOVA with Tukey’s multiple comparison test. **(e)** Schematic of *in vivo* Cre lentivirus delivery through nasal instillation, resulting in recombination and YFP expression in the mouse lung. **(f)** GFP expression in mouse lung parenchyma 10 days (middle) and 4 weeks (right) following nasal instillation of CMV-GFP lentivirus, and untreated control (left), scale bars 100 μm. **(g)** Varying degrees of Cre-mediated recombination and YFP expression (low to high) in R26(LSL-YFP) reporter mouse lung parenchyma 10 days following nasal instillation with CMV-Cre lentivirus, scale bars 100 μm. **(h)** IHC quantification of Cre-mediated YFP recombination (YFP+ cells per mm2) in R26(LSL-YFP) reporter mice 10 days following nasal instillation with varying doses of CMV-Cre lentivirus. Error bars indicate standard error of the mean. **(i)** Comparison of the rate of CMV-Cre and Ef1a-Cre driven recombination and YFP expression in R26(LSL-YFP) reporter mice, 10 days following nasal instillation of Cre-lentivirus, unpaired, two-tailed Student’s t-test. **(j)** PCR analysis showing retained viral payload (Cre sequence, left), and YFP (middle) and Kras (right) recombination in lung tumor DNA from CMV-Cre transduction-initiated tumors in P53(F/F), Kras(LSL-KrasG12D/+), Rosa26(LSL-YFP); PPKY mice, with somatic (tail) DNA from the same mouse as control. **(k)** IHC analysis revealing sporadic Cre expression in lung parenchymal 10 days following nasal instillation of CMV-Cre lentivirus (left) and in lentivirus transduction-initiated lung tumor (P53(F/F), Kras(LSL-G12D/+), R26(LSL/YFP); PPKY/LV) at survival endpoint (right), scale bar 100 μm.

In summary, following successful high titer lentivirus production, reliable titer quantification, critical comparison of promoter and dose dependencies as well as the confirmation of persistent gene expression, we were able generate lung tumors with stable genomic integration.

### Oncogene activation accelerates transduction-initiated lung tumor progression

We next set out to test if we can trigger CRISPRa-mediated oncogene activation and tumor formation *in vivo*. To generate CRISPRa lung tumors, we administered 1e8TU Cre/sgMyc or Cre/sgYap1 encoding lentivirus through nasal instillation to PPKS mice (referred to as PPKS/M and PPKS/Y, respectively) (Figure 4a). In addition, we administered lentivirus encoding for Cre and a non-targeting guide to PPKS (PPKS/NT) and PPKY (PPKY/LV) mice to serve as non-targeted CRISPRa and non-CRISPRa controls, respectively (Supplementary Table S3).

**Figure 4.**
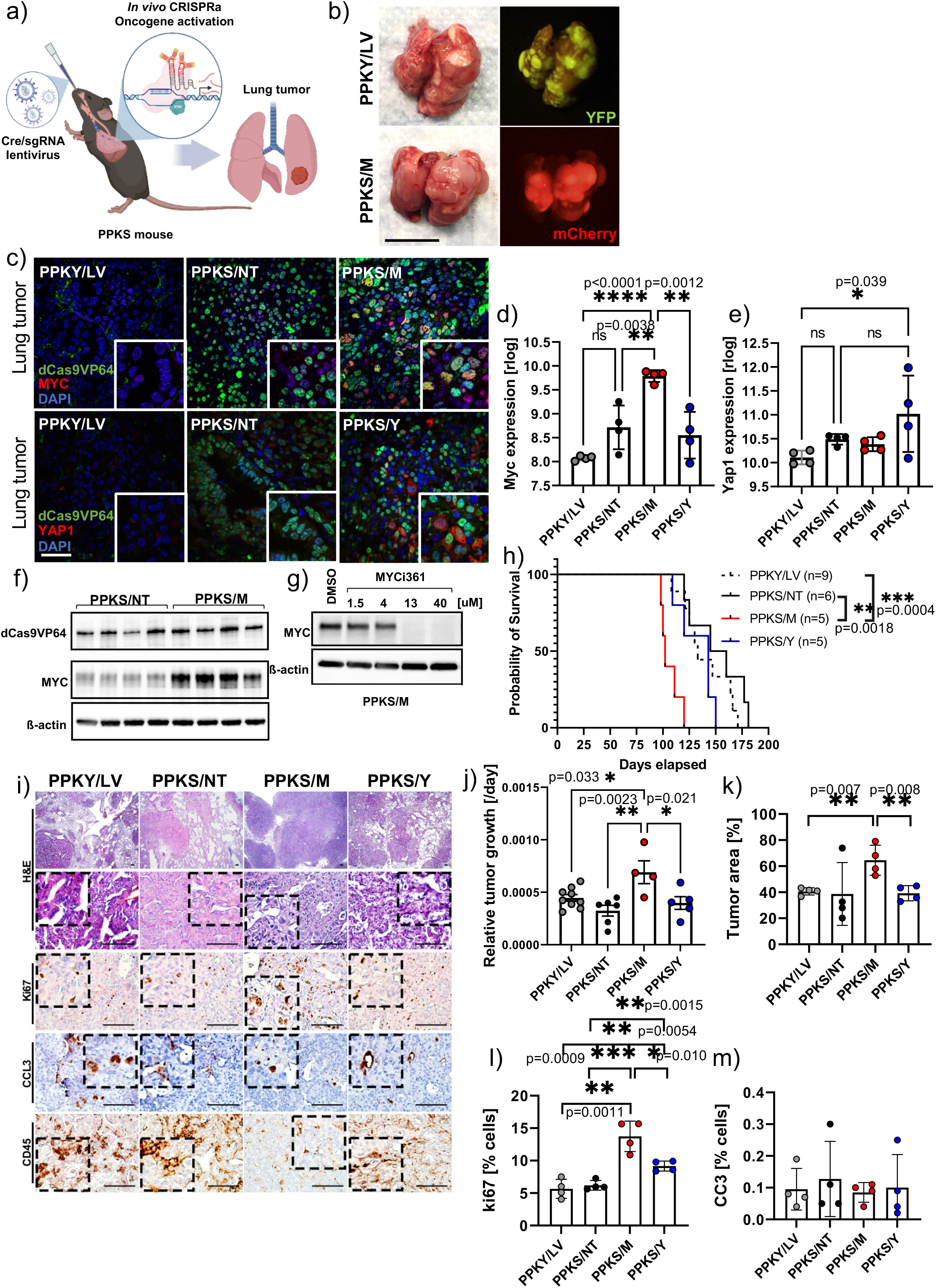
Tumor-initiation and *in vivo* oncogene activation through lentiviral transduction in the CRISPRa/SAM lung tumor model. **(a)** Schematic of the autochthonous CRISPRa/SAM gene activation lung tumor model. Programmable oncogene activation and tumor-initiation is triggered through nasal instillation of oncogene-targeted Cre/sgRNA lentivirus (CMV-Cre-U6-sgRNA(MS2)) in CRISPRa-competent P53(F/F), Kras(LSL-KrasG12D/+), Rosa26(LSL-SAM); PPKS mice. **(b)** Macroscopic and fluorescent images of CMV-Cre-U6-sgRNA(MS2) lentivirus-induced lung tumors in non-CRISPRa P53(F/F), Kras(LSL-KrasG12D/+), Rosa26(LSL-YFP); PPKY (top), and CRISPRa-competent PPKS mice (bottom), expressing YFP and mCherry, respectively, scale bar 1cm. **(c)** Immunofluorescent analysis of Myc-activated (PPKS/M) and Yap1-activated (PPKS/Y) lung tumors, with non-CRISPRa (PPKY/LV) and non-targeted CRISPRa (PPKS/NT) controls. All PPKS tumor reveal nuclear dCas9VP64 expression, while increased nuclear MYC and predominantly cytoplasmic YAP1-expression was detected in PPKS/M and PPKS/Y tumors, respectively. dCas9VP64 (green), MYC/YAP1 (red), DAPI (blue). Scale bar 40 μm. **(d)** RNA-seq analysis revealing *Myc* over-expression in *Myc*-activated (PPKS/M) tumors relative to controls, *Myc* expression levels displayed as rlog values, one-way ANOVA with Tukey’s multiple comparison test. **(e)** RNA-seq analysis reveals *Yap1* over-expression in *Yap1*-activated (PPKS/Y) tumors relative to controls, *Yap1* expression displayed as rlog values, one-way ANOVA with Tukey’s multiple comparison test. **(f)** Immunoblot analysis of dCas9VP64, MYC and ß-actin in cell lines derived from *Myc*-activated (PPKS/M) and non-targeted control (PPKS/NT) lung tumors revealing MYC-overexpression in PPKS/M tumor-derived cell lines relative to controls and retained dCas9VP64 expression in all cell lines. **(g)** Immunoblot of MYC and ß-actin in mouse tumor cells derived from *Myc*-activated lung tumors (PPKS/M) treated with various concentrations of MYC-inhibitor (MYCi361). **(h)** Kaplan-Meier survival analysis of CMV-Cre-U6-sgRNA(MS2) lentivirus induced lung tumors in PPKY and PPKS mice, revealing significantly shorter survival for *Myc*-activated (PPKS/M) tumor mice relative to non-CRISPRa (PPKY/LV) and non-targeted CRISPRa (PPKS/NT) control tumor mice. Pairwise Mantel-Cox (logrank) test. **(i)** Representative tumor histology (HE), ki67, cleaved Caspase 3 (CC3) and CD45 staining in *Myc*-activated (PPKS/M) and *Yap1*-activated (PPKS/Y) mouse lung tumors, next to non-CRISPRa (PPKY/LV) and non-targeted CRISPRa (PPKS/NT) control tumors, scale bar 100 μm. **(j)** Relative tumor growth rate (tumor weight/(body weight*survival time) for each cohort, one-way ANOVA with Tukey’s multiple comparison test. **(k)** Quantification of relative tumor area reveals increased tumor burden in *Myc*-activated (PPKS/M) tumors relative to PPKY/LV and PPKS/NT controls. **(l)** IHC analysis of ki67 expression reveals increased proliferation rate in *Myc*-(PPKS/M) and *Yap1*-activated (PPKS/Y) lung tumors relative to PPKY/LV and PPKS/NT controls. **(m)** IHC analysis of cleaved Caspase 3 (CC3) expression revealed no significant differences in apoptotic rate between tumor cohorts. **(k), (l),(m)** significance tested with two-tailed Student’s t-test

All mice treated with Cre lentivirus through nasal instillation developed mCherry or YFP-positive lung tumors (Figure 4b). To determine if oncogene activation led to persistent *Myc* and *Yap1* overexpression in the resulting tumors, we stained tumor sections for MYC, YAP1 and dCas9VP64, and performed mRNA-seq on tumor tissues (Figures 4c-e). Immunofluorescent staining revealed marked nuclear MYC expression in PPKS/M tumors and predominantly cytoplasmic YAP1 expression in PPKS/Y tumors, relative to controls (Figure 4c). Transcriptomic analysis revealed that *Myc* was overexpressed 2.8 and 6.5-fold in *Myc*-activated (PPKS/M) tumors, relative to non-targeted (PPKS/NT) and non-CRISPRa (PPKY/LV) tumors, respectively (Figure 4d). *Yap1* was significantly overexpressed 3.3-fold in *Yap1*-activated (PPKS/Y) tumors relative to non-CRISPRa (PPKY/LV) tumors. We found 2.2-fold overexpression of *Yap1* relative to non-targeted tumors (PPKS/NT), but the result did not reach statistical significance (Figure 4e). In addition, we confirmed sustained MYC over-expression in cell lines derived from PPKS/M tumors, relative to PPKS/NT controls, and retained dCas9VP64 expression in all PPKS cell lines tested (Figure 4f, Supplemental Figure 4a-b). Using a recently described MYC-inhibitor (MYCi361(21)), we were able to pharmacologically suppress MYC in these cell lines (Figure 4g).

Strikingly, *Myc* activation resulted in significantly decreased overall survival (median survival: PPKS/M 102 days, PPKS/NT 152.5 days, PPKY/LV 133 days, Figure 4h, Supplementary Table S3)), increased tumor burden (Figure 4i-k) and higher proliferation rates (Figure 4i,l) relative to controls, while apoptosis remained unaffected (Figure 4i,m). Although an increase in proliferation was observed in PPKS/Y relative to controls (Figure 4l), no significant difference in tumor progression or survival was obvious between PPKS/Y and PPKS/NT or PPKY/LV controls, nor between PPKS/NT and PPKY/LV (Figure 4h), indicating a specific oncogenic effect of *Myc*, but not *Yap1*, activation in the lung.

### *Myc*-activation drives an aggressive, immune-suppressed tumor phenotype

As *Myc* activation resulted in markedly more aggressive tumor progression, indicated by decreased overall survival, increased tumor burden and tumor cell proliferation, we analyzed the transcriptomic features and phenotype of these tumors in more detail.

To investigate the effect of *Myc* activation on MYC network signaling, we analyzed the expression of the main *Myc* family members (*Myc*, *Mycn* and *Mycl*) and the proximal MYC signaling network (Supplementary Table S4,(23)) in *Myc*-activated (PPKS/M) relative to non-targeted control (PPKS/NT) tumors. *Myc*-activated tumors displayed significant over-expression of *Mxi1*, *Mlx* and *Mxd3* in addition to *Myc*, whereas *Mycn*, *Mycl*, *Mxd1*, *Mlxip*, *Mnt* and *Mxd4* were significantly down-regulated, indicating a pronounced effect on the proximal MYC signaling network (Figure 5a). Down-regulation of *Mycn* and *Mycl* in *Myc*-activated tumors may represent a compensatory mechanism in response to *Myc*-overexpression.

**Figure 5.**
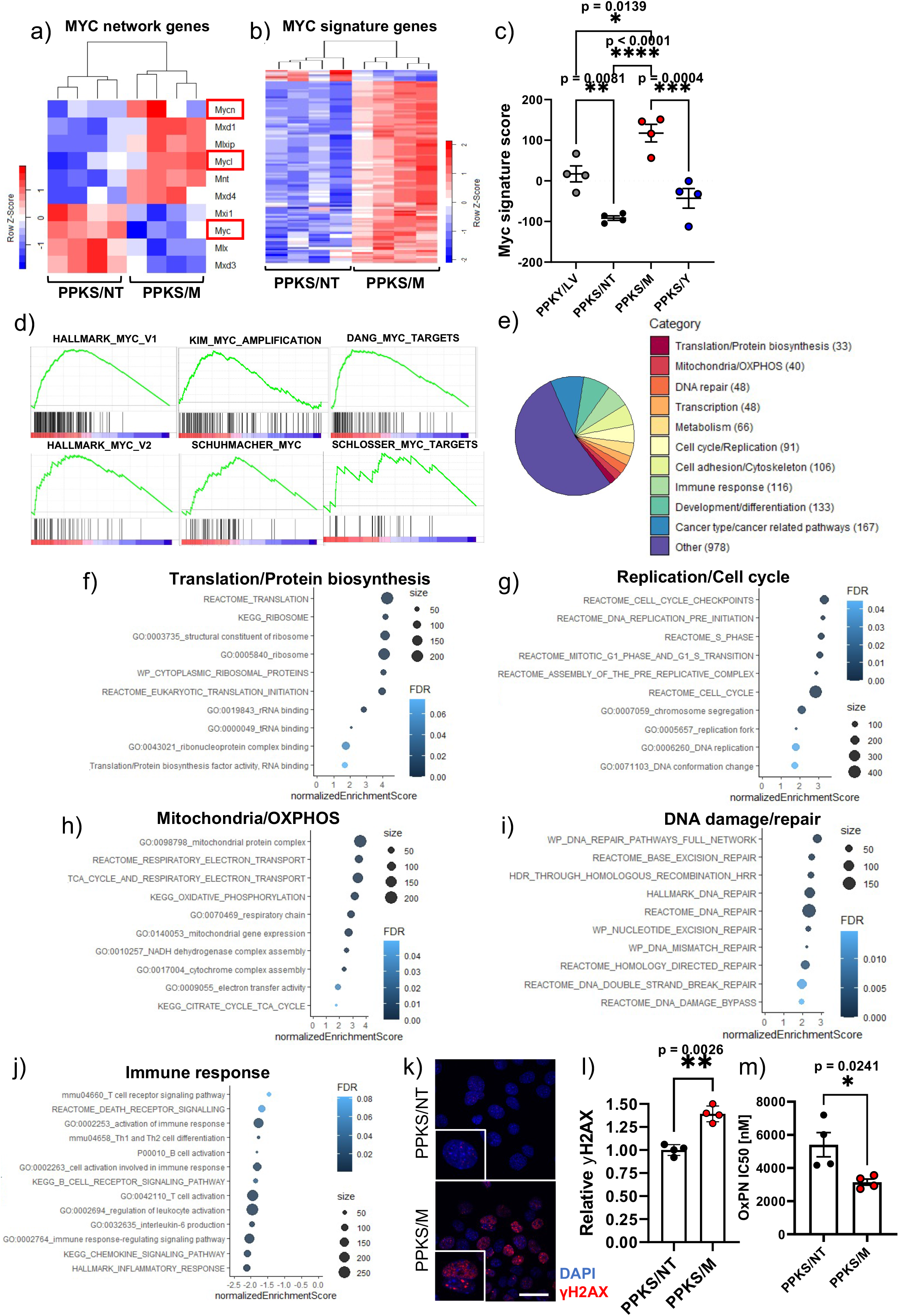
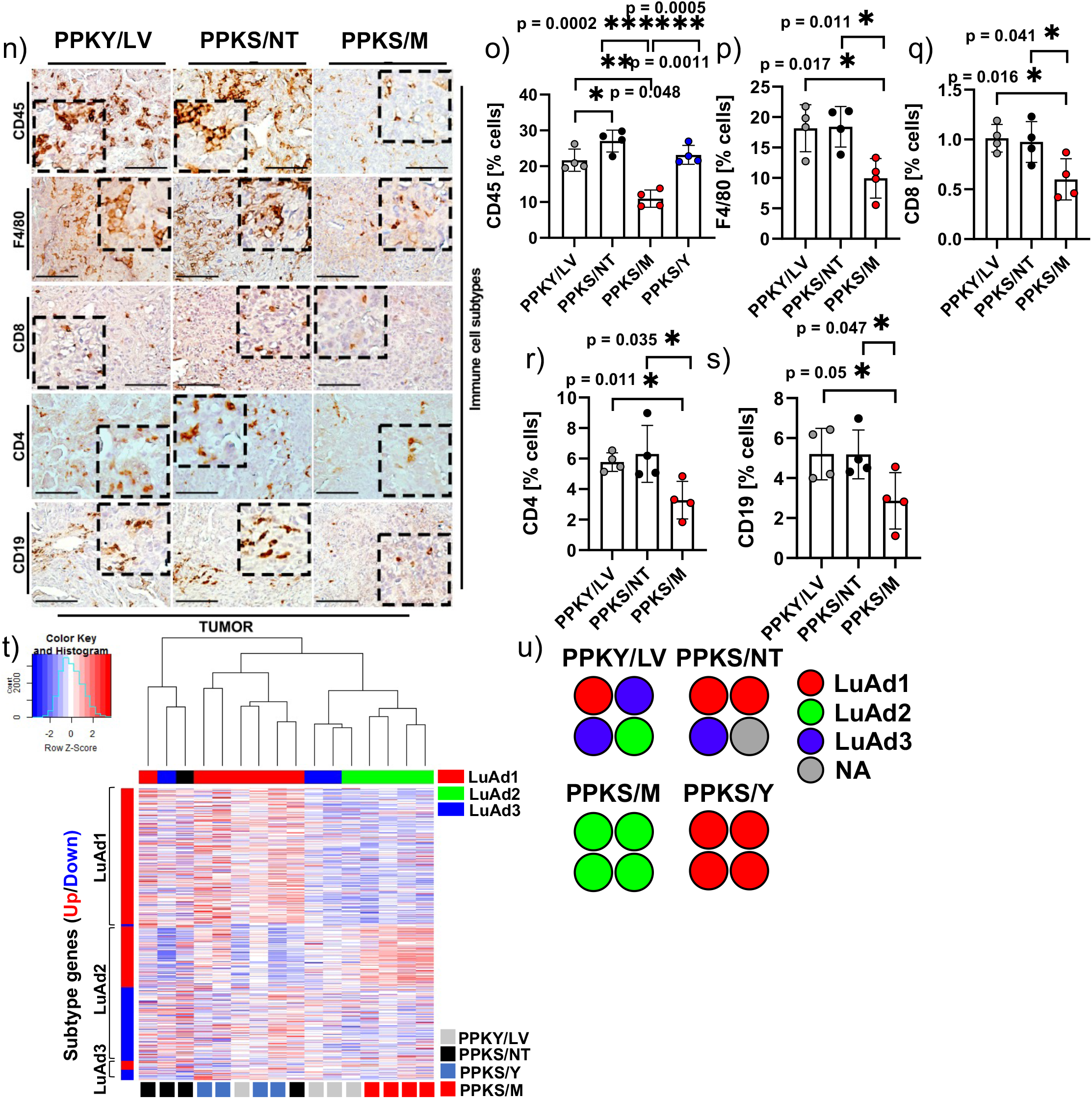
Transcriptomic reprogramming in *Myc*-activated lung tumors from the CRISPRa/SAM model. **(a)** Heatmap of differentially expressed MYC signaling network members in *Myc*-activated (PPKS/M) tumors relative to non-targeted (PPKS/NT) tumor controls, MYC/N/L marked in red. **(b)** Heatmap showing differential expression of 107 MYC-signature genes in PPKS/M tumors relative to PPKS/NT controls, 100 genes displaying over-expression and 7 displaying down-regulation. **(c)** MYC signature score calcuated from 134 MYC signature genes, one-way ANOVA with Tukey’s multiple comparison test, **(d)** Gene set enrichment analysis (GSEA) revealing positive enrichment of MYC-associated gene sets in *Myc*-activated (PPKS/M) lung tumors relative to non-targeted (PPKS/NT) tumor controls. **(e)** GSEA analysis revealing enrichment of a large number of gene sets associated with canonical MYC function. **(f)** Enrichment dot-plot of a selection of translation/protein biosynthesis associated gene sets enriched in *Myc*-activated (PPKS/M) relative to non-targeted (PPKS/NT) tumors. **(g)** Enrichment dot-plot of a selection of replication/cell cycle associated gene sets enriched in *Myc*-activated (PPKS/M) relative to non-targeted (PPKS/NT) tumors. **(h)** Enrichment dot-plot of a selection of mitochondria/oxidative phosphorylation (OXPHOS) associated gene sets enriched in *Myc*-activated (PPKS/M) relative to non-targeted (PPKS/NT) tumors. **(i)** Enrichment dot-plot of a selection of DNA damage/repair associated gene sets enriched in *Myc*-activated (PPKS/M) relative to non-targeted (PPKS/NT) tumors. **(j)** Enrichment dot-plot of a selection of immune response associated gene sets negatively enriched in *Myc*-activated (PPKS/M) relative to non-targeted (PPKS/NT) tumors. **(k)** Immunofluorescent analysis of γH2AX foci in in cells derived from *Myc*-activated (PPKS/M) tumors and non-targeted controls (PPKS/NT), γH2AX (red), DAPI (blue), scale bar 20um. **(l)** Quantitative analysis of γH2AX foci reveals increased DNA damage in cells derived from *Myc*-activated (PPKS/M) tumors, relative to non-targeted controls (PPKS/NT). Two-tailed Student’s t-test **(m)** Drug response analysis reveals that cells derived from *Myc*-activated (PPKS/M) lung tumors are significantly more sensitive to oxaliplatin than cells derived from non-targeted control (PPKS/NT) tumors, as indicated by lower IC50 values, two-tailed Student’s t-test. **(n)** IHC analysis of CD45+ immune cells, F4/80+ macrophages, CD8+ T cells, CD4+ T cells and CD19+ B cells in *Myc*-activated (PPKS/M) and control (PPKY/LV and PPKS/NT) tumors, scale bar 100 μm. **(o)** Quantitative IHC of CD45+ immune cells. **(p)** Quantitative IHC analysis of F4/80+ macrophages. **(q)** Quantitative IHC analysis of CD8+ T cells. **(r)** Quantitative IHC analysis of CD4+ T. **(s)** Quantitative IHC analysis of CD19+ B cells. **(t)** Heatmap and unsupervised hierarchical clustering of mouse lung tumors based on expression of transcriptomic subtyping genes, and CMScaller classification based on over-expressed subtype-specific genes. **(u)** CMScaller subtype classification of mouse tumors by tumor cohort. **(o)**, **(p)**, **(q)**, **(r)**, **(s)** significance tested with two-tailed Student’s t-test

To infer the effect of *Myc*-activation on the transcription of down-stream targets we probed the expression of a list of 134 MYC-signature genes with evidence of direct *Myc*-binding (see Supplementary Table S4,(24–26)). We found that 100 of these genes (75%) were significantly over-expressed in *Myc*-activated tumors (PPKS/M) relative to non-targeted controls (PPKS/NT), whereas 7 genes (5%) were down-regulated (Figure 5b). Using the expression values for these genes, we calculated a MYC signature score (see methods) for each tumor which was significantly higher in PPKS/M tumors compared to all other groups (Figure 5c).

To determine if we could detect a *Myc*-driven tumor phenotype, we analyzed our model for known transcriptomic *Myc*-associated gene sets by performing Gene Set Enrichment Analysis on genes differentially expressed in PPKS/M tumors relative to non-targeted controls (PPKS/NT). GSEA revealed strong enrichment for multiple sets of MYC target genes (Figure 5d). In addition, GSEA showed strong positive enrichment for canonical MYC-associated biological processes in *Myc*-activated tumors, including, translation/protein biosynthesis, replication/cell cycle, DNA damage response/DNA repair and oxidative phosphorylation/mitochondrial function (Figure 5e-i, Supplementary Table S5). Importantly, we found that a large number of immune response-related pathways were negatively enriched (Figure 5j). To confirm some of these findings *in vitro*, we next set out to determine if cancer cells from *Myc*-activated tumors displayed an objective increase in baseline DNA damage. Indeed, we found that cell lines derived from PPKS/M tumors displayed significantly increased baseline DNA damage compared to controls, as indicated by an increase in nuclear γH2AX foci (Figure 5k-l). We also found that PPKS/M cells were significantly more sensitive to oxaliplatin relative to PPKS/NT cells, as indicated by lower oxaliplatin IC50 values, 3.4uM and 5.4uM, respectively (Figure 5m)

There is accumulating evidence that *Myc* expression in non-small cell lung cancer (NSCLC) and other tumor entities drives an immune suppressive microenvironment(18–21,27), which is in line with our GSEA findings of negative enrichment of numerous immune related pathways (Figure 5j). To investigate this potential link, we set out to decipher the immune composition in our PPKS/M lung tumors compared to controls, using IHC-based immune subtyping. Next to a significant overall decrease in immune cells (CD45^+^ cells) in *Myc* activated tumors, we found that the population of cytotoxic CD8^+^ T cells and CD4^+^ T helper-cells, B-cells (CD19^+^) and macrophages (F4/80^+^) was significantly decreased compared to controls (Figure 5n-s). These observations reveal a profound effect of *Myc* activation in driving a less immunogenic, or “immune cold”, and likely immune suppressive tumor microenvironment in our model. Taken together, these results show that our model is able to faithfully recapitulate numerous MYC-associated features and serves to indicate the wide-reaching potential of our approach.

### Transcriptomic subtyping of murine and human lung adenocarcinomas reveals MYC as a driver of the LuAd2 subtype

Recently, Jang et al. classified a cohort of human lung adenocarcinoma tumors into three subtypes (LuAd1-3) with distinct transcriptomic and immune-regulatory features, and potential immunotherapy-response predictive power(22). LuAd1 was found to represent an “immune-hot” tumor subtype with potentially favorable response to immune therapy, whereas LuAd2 and LuAd3 represented distinct “immune-cold” tumor subtypes (Supplementary Table S6). However, the transcriptomic drivers of these subtypes remain to be elucidated. To characterize our mouse tumors using an analogous approach, we subtyped the tumors with CMScaller(28) using lists of subtype-specific, over-expressed gene orthologs from the Jang et al. study. Tumor subtyping revealed starkly different transcriptomic profiles between individual mouse tumors (Figure 5t, Supplementary Table S6). While all *Yap1*-activated tumors were subtyped as LuAd1, strikingly, all PPKS/M tumors (4 out of 4) were subtyped as LuAd2 and clustered separately from the other tumors (Figure 5t-u). Based on this observation we hypothesized that *Myc* is a potential driver of the LuAd2 subtype.

To expand this analysis and to investigate a potential link between MYC signaling and tumor subtype in human lung adenocarcinomas, we applied a similar approach to stratify the 585 lung adenocarcinoma cases in the TCGA-LUAD cohort. In summary, 206 (35%), 256 (44%) and 93 (16%) patient tumors were subtyped as LuAd1, LuAd2 and LuAd3, respectively, while 30 (5%) patient tumors did not classify as any subtype (Figure 6a, Supplemental Figure 5a, Supplementary Table S6).

**Figure 6.**
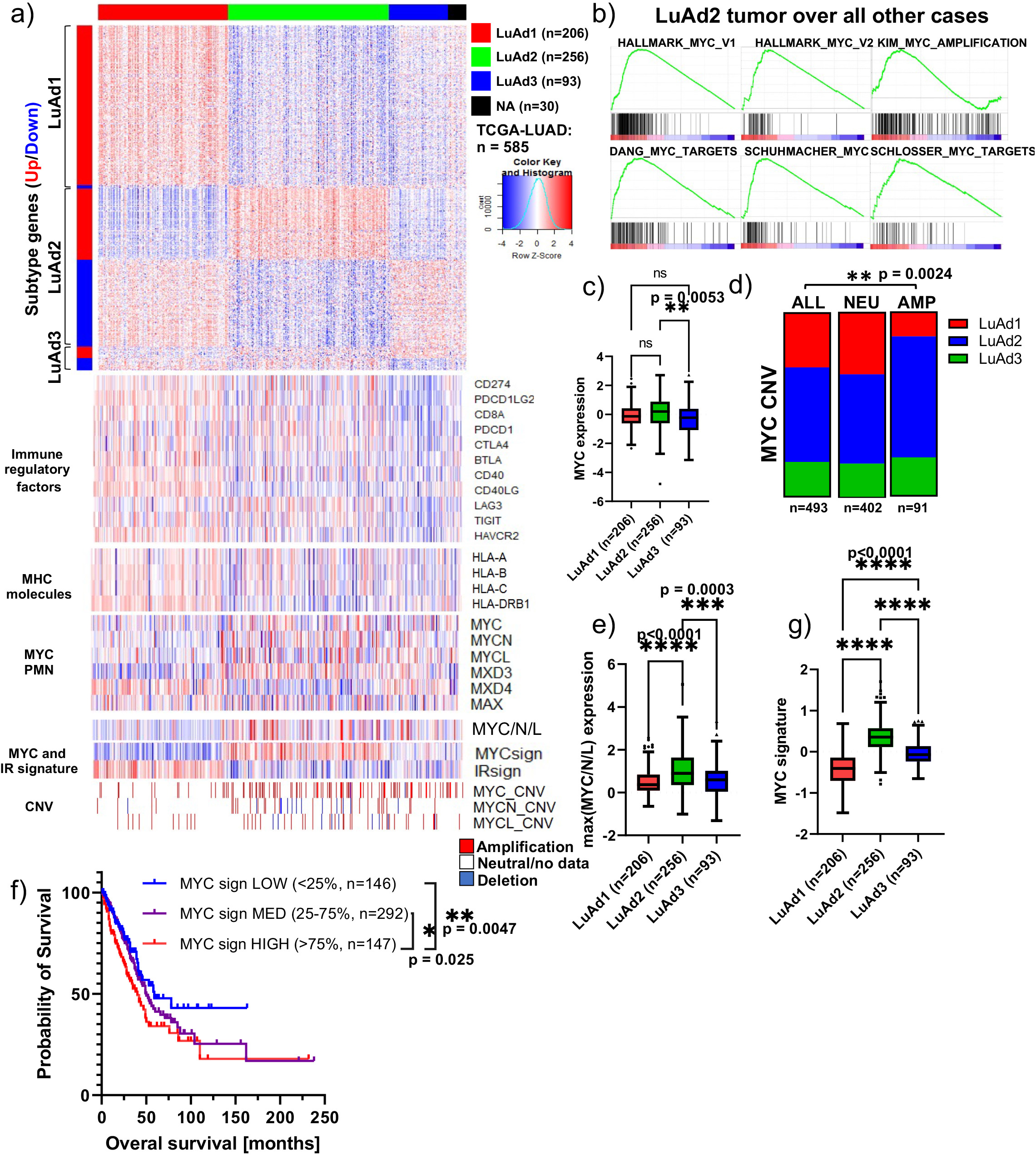
Transcriptomic subtyping of lung adenocarcinoma tumors reveals enrichment of MYC signature in LuAd2 patient tumors. **(a)** Heatmap of subtype-specific gene expression in TCGA-LUAD patient tumors organized by CMScaller subtype classification (LuAd1/LuAd2/LuAd3), and heatmaps of selected immune regulatory genes, MHC molecules, MYC signaling network members, maximum MYC/N/L expression, MYC signature, IR signature, and *MYC*, *MYCN* and *MYCL* copy number variation. **(b)** Gene set enrichment analysis revealing positive enrichment for several MYC-associated gene sets in LuAd2 tumors relative to all other LuAd tumors. **(c)** Normalized *MYC* expression in LuAd1, LuAd2 and LuAd3 tumors from the TCGA-LUAD data set, one-way ANOVA with Tukey’s multiple comparison test. **(d)** LUAD subtype distribution as function of *MYC* copy number status revealing enrichment of LuAd2 and depletion of LuAd1 among *MYC* amplified tumors, chi-squared test. **(e)** Maximum normalized expression of *MYC*, *MYCN* or *MYCL* from each tumor as function of tumor subtype, one-way ANOVA with Tukey’s multiple comparison test. **(f)** Kaplan-Meier survival plot of the TCGA-LUAD data set, stratified by patient tumor MYC-signature; MYC-signature LOW (<25 percentile, n=146), MED (25-75 percentile, n=292), HIGH (>75 percentile, n=147), pairwise Mantel-Cox (logrank) test. **(g)** MYC signature score in LuAd1, LuAd2 and LuAd3 tumors from the TCGA-LUAD data set, one-way ANOVA with Tukey’s multiple comparison test.

Interestingly, we found that multiple MYC-associated gene sets were strongly enriched in LuAd2 tumors over all other cases (Figure 6b). We found that LuAd2 tumors expressed significantly more MYC relative to LuAd3, and MYC copy-number amplified tumors were significantly enriched for the LuAd2 subtype (Figure 6a,c-d). When we considered the maximum expression value for *MYC*, *MYCN* and *MYCL* for each tumor, we found that LuAd2 expressed significantly higher maximum MYC/N/L levels compared both other groups (Figure 6a,e), consistent with a direct link between the MYC/MYCN/MYCL signaling axis and LuAd2.

Next, we defined a MYC signature score based on the expression of 132 well-established MYC-target genes(24–26). Stratification of the TCGA-cohort into 3 cohorts, revealed that a high MYC signature score was associated with significantly shorter survival relative to MYC signature medium and low tumors (median survival 40 (high), 50 (medium) and 59 months (low), Figure 6f). Consistent with our hypothesis, we found that LuAd2 tumors displayed a strongly elevated MYC signature score as compared to LuAd1 and LuAd3 (Figure 6a,g). Taken together, transcriptomic subtyping of tumors obtained from our CRISPRa/SAM Myc activation mouse model and human lung adenocarcinoma cohorts (TCGA) allowed us to identify MYC as a potential driver for the LuAd2 subtype, warranting further investigation of the link between MYC and tumor immune landscape.

### MYC-signaling as a driver of an immune-suppressive lung adenocarcinoma subtype 2 may predict immune therapy response

Consistent with Jang et al., we found that LuAd2 and LuAd3 tumors in the TCGA-LUAD data set express significantly lower levels of a set of immune regulatory factors, and MHC molecules relative to LuAd1 tumors (Figure 6a, Figure 7a-b), consistent with these subtypes representing “immune-cold” subtypes and LuAd1 representing an “immune-hot” subtype. Furthermore, we found that LuAd2 and LuAd3 tumors displayed significantly lower immune response score (IRscore)(22) relative to LuAd1 (Figure 7c, Supplementary Table S6). Moreover, the LuAd2 subtype was associated with significantly shorter survival as compared to LuAd1 (median survival 38.47 and 71.42 months, p<0.0001, Figure 7d), while no significant difference in survival was observed between LuAd2 and LuAd3 nor between LuAd3 and LuAd1. We were thus able to validate the major findings from the Jang et al. study across a much larger cohort of patient tumors.

**Figure 7.**
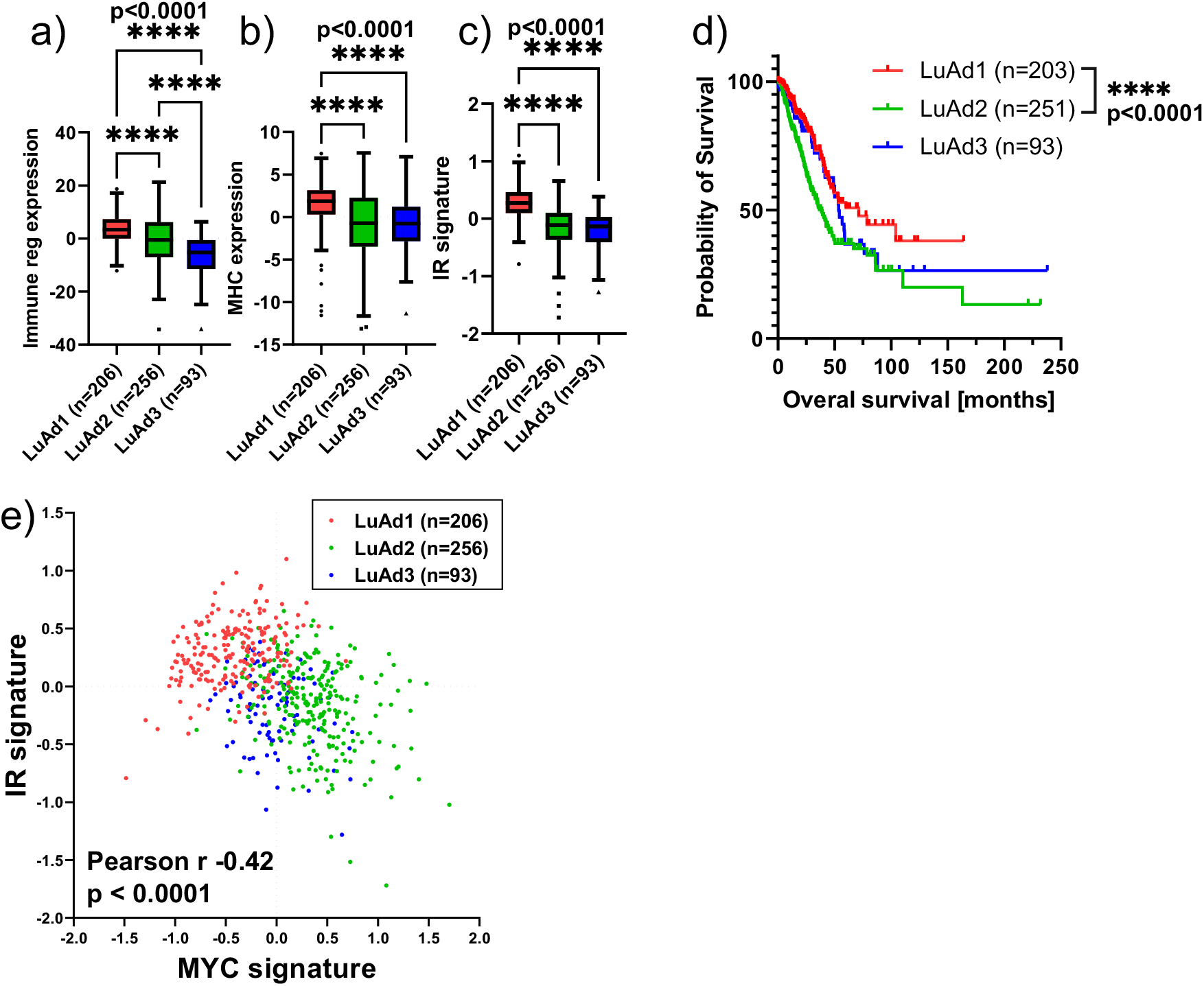
MYC signature and immune response. **(a)** Normalized expression of immune regulatory factors as function of LuAd subtype in the TCGA-LUAD data set, one-way ANOVA with Tukey’s multiple comparison test. **(b)** Normalized expression of MHC factors as function of LuAd subtype in the TCGA-LUAD data set, one-way ANOVA with Tukey’s multiple comparison test. **(c)** IR-signature as function of LuAd subtype in the TCGA-LUAD data set, one-way ANOVA with Tukey’s multiple comparison test. **(d)** Kaplan-Meier survival plot of the TCGA-LUAD data set, stratified by patient tumor subtype; LuAd1 (n=203), LuAd2 (n=251) and LuAd3 (n=93). Pairwise Mantel-Cox (logrank) test. **(e)** Correlation plot of IR signature and MYC signature in patient tumors from the TCGA-LUAD data set, Pearson’s correlation coefficient (r= −0.42, p<0.0001).

Furthermore, we found that the IRscore was significantly negatively correlated with the MYC signature score across the TCGA-LUAD cohort (Pearson r = −0.42, p<0.0001), consistent with an antagonistic relationship between MYC signaling and factors important for response to immune therapy (Figure 7e). We also implemented our analysis on an independent dataset from OncoSG (169 patients). Within this data set we were able to recapitulate our main findings from the TCGA dataset; LuAd2 tumors displayed significantly higher MYC signature scores and lower IRscores as compared to LuAd1, and the MYC signature score again predicted survival. (Supplemental Figure 5b–1)

Overall, these results, combined with the results from our mouse model, strongly implicate MYC signaling as a driver of the “immune cold” LuAd2 subtype. This highlights the potential of MYC-targeted pharmacological intervention to improve patient responses to immune checkpoint blockade.

## Discussion

Dysregulation of gene expression is a key characteristic of many human diseases, including cancer. Introducing controlled gene perturbations in animal models through genetic engineering is necessary for creating models that faithfully recapitulate human disease. The groundbreaking developments in genetic engineering of the last decade have opened new avenues for more flexible and sophisticated models. Here, we have demonstrated that our CRISPRa/LSL-SAM mouse model is a powerful and versatile tool for programmable gene activation *in vitro* and *in vivo*, and we have demonstrated its use to investigate oncogene drivers in cancer and tumor formation. We have shown that gene activation can be triggered in primary cells *ex vivo*, and that the resulting cells can be transplanted into syngeneic mice to generate tumors. In addition to allowing for the study or individual genes, this approach could straightforwardly be expanded to *in vitro* or *in vivo* screening in various cell types using pools of gene-targeting guides. Importantly, we detected only moderate levels of transcriptional gene activation (2.8 – 4.4 fold over-expression), i.e. expression levels only marginally supraphysiological. This is in stark contrast to conventional, viral promoter driven/transgene approaches where over-expression can reach hundred-fold, potentially resulting in artificial rather than physiologically relevant effects We have also demonstrated that gene activation can be triggered directly *in vivo* through lentiviral transduction and that oncogene activation in this model can lead to dramatic differences in tumor characteristics and phenotype, recapitulating observations from human patients. For this purpose, we developed reliable protocols for in-house reproducible customized high titer production, titer quantification, and determined promoter specificity and transfections efficiencies. The direct *in vivo* transduction approach can likely also be adapted to allow for limited-scale oncogene screening through the use of pooled lentiviral guides, efforts which we are currently pursuing.

By activating the Myc oncogene concurrently with oncogenic Kras and P53 loss, we were able to recapitulate important MYC-associated features of human tumors in our model. The results from our model also generated the hypothesis of a direct link between MYC signaling, an immune suppressive tumor microenvironment and a clinically relevant and transcriptomically defined tumor subtype in lung adenocarcinoma (LuAd2). Through cross-species validation, our present work highlighted a significant cohort of human lung adenocarcinoma patient tumors transcriptomically defined as subtype 2 and characterized by a pronounced Myc signaling signature and immune suppression. This MYC-signature high/LuAd2 subtype patient cohort may display enhanced response to combined MYC/immune-targeting therapies.

## Methods

### Generating the LSL-SAM mouse

To target conditional dCas9VP64, MS2-p65-HSF1 and mCherry expression to the Rosa26 locus in embryonic mouse stem cells (mES), we generated the pLSL-dCas9VP64-P2A-MS2-p65-HSF1-T2A-mCherry-Rosa26TV (LSL-SAM) vector: a 6.7kbp fragment encoding for tricistronic expression of dCas9VP64, MS2-p65-HSF1 and mCherry was amplified from the PB-UniSAM vector and assembled into the BstBI/NsiI digested empty backbone of the pLSL-Cas9-Rosa26TV vector, using HiFi DNA assembly (NEB). The resulting Rosa26 targeting vector encodes for conditional, CAG-promoter driven dCas9VP64, MS2-p65-HSF1 and mCherry expression, incorporating positive (G418) and negative (DTA) selection. Stbl3 bacterial colonies were screened for the correct construct with BstBI/NsiI digestion and agarose electrophoresis. The LSL-SAM insert from one bacterial colony was sequence-verified trough NGS denovo assembly (Applied Biological Materials Inc.), linearized through XhoI digestion and electroporated into G4 mES cells. Following selection with G418, approximately 200 mES clones were screened for successful Rosa26 targeting using PCR reactions with primers binding inside and outside the right and left homology arms, with an overall success rate of approximately 60%. Two clones were expanded for blastocyst injection and chimera generation through implantation in albino B6 females. Founder mice with germline transmission were generated from both mES clones, and one clone was used for all further experiments. Electroporation, selection, colony picking and blastocyst implantation was performed by the MD Anderson Cancer Center Genetically Engineered Mouse Facility (GEMF). LSL-SAM mice were genotyped using a 3-primer PCR reaction that recognizes the R26(LSL-SAM) construct as well as the wild type Rosa26 allele, see Supplementary table S7 for genotyping primers.

#### Mouse models

All studies were conducted in compliance with the institutional guidelines (MD Anderson; IACUC). The following mouse models were used in these studies: *Rosa26*^*LSL-SAM*^ (LSL-SAM), *Rosa26*^*LSL-YFP*^ (LSL-YFP, Rhim 2012), *P53*^*FlF*^*;Kras*^*LSL-G12D/+*^*;Rosa26*^*LSL-YFP*^ (PPKY(29)), *P53*^*FlF*^*;Kras*^*LSL-G12D/+*^*; Rosa26*^*LSL-SAM*^ (PPKS). Mice were backcrossed to a C57BL/6NJ background (see Supplementary Table S8). All animals were archived, and mice with the same genotypes were assigned randomly to experimental groups. All experiments were performed on balanced cohorts of male and female mice. We found no sex-specific differences in disease initiation and progression. All mice with the appropriate genotype were included in the study. All mice received standard chow diet *ad libitum* and were housed in pathogen-free facility (modified barrier) with standard controlled conditions (temperature 21 to 23°C; set point 22°C; relative humidity: 40-55%; set point 45%). No more than 5 mice were housed together, under the supervision of DVMS veterinarians, in an AALAC-accredited animal facility at the University of Texas M.D. Anderson Cancer Center. The number of mice used for each experiment are listed as (n). All animal procedures were reviewed and approved by the MDACC Institutional Animal Care and Use Committee (IACUC 00001626, PI: Rhim). Maximal tumor burden (10% of body weight) and maximal tumor size allowed by the ethics committee was not exceeded. No statistical methods were used to pre-determine sample sizes for *in vivo* experiments, but our sample sizes are similar to those reported in previous publications (30).

#### Cell lines

Lenti-X 293T cell line (Takara; #632180) and tumor-derived murine cell lines were cultured in DMEM with 10% FBS with 1x penicillin/streptomycin at 37°C in a humidified incubator with 5% CO2 and tested regularly for mycoplasma (see Supplementary Table S9).

### Copy Number Assay

LSL-SAM copy number was determined using the TaqMan Copy Number Assay following the manufacturer’s instructions using a custom mCherry TaqMan assay and Tfrc as the reference gene. Mice with known single mCherry copy number were used as controls.

### Cre/CRISPRa backbone vector cloning

The pLV-Ef1a-Cre-U6-sgRNA(MS2) vector was generated by replacing the BleoR cassette with Cre recombinase from the pBS513 EF1alpha-Cre vector in the plenti sgRNA(MS2)_zeo backbone using PCR and HiFi assembly (NEB). One BsmBI site in the Cre cassette was destroyed through targeted mutagenesis. The pLV-CMV-Cre-U6-sgRNA(MS2) vector was generated by replacing the Ef1a promoter with a CMV promoter cassette in pLV-Ef1a-Cre-U6-sgRNA(MS2) using PCR and HiFi assembly (NEB). The pLV-CMV-CreERT2-P2A-[puroR/zeoR/hygR]-U6-sgRNA(MS2) vectors were generated by replacing the Ef1a-puroR cassette in plenti_sgRNA(MS2)_puro with CMV-CreERT2-P2A-[puroR/zeoR/hygR] cassettes derived from pMXS_CMVCreERT2-bGHpA, pLCE-P2A-Puro, plenty_sgRNA(MS2)_zeo and pLenti CMV rtTA3 Hygro. Two BsmBI sites in the CreERT2 cassette were destroyed through targeted mutagenesis. All generated plasmids were sequence verified with Sanger sequencing (see Supplementary Table S10). All reagents are listed in Supplementary Table S11.

### Cre/CRISPRa guide cloning

sgRNA sequences were cloned into the Cre-encoding lentiviral backbones with Golden Gate cloning using a previously described protocol(8). Briefly, 100 pmol of complementary sgRNA sequence containing DNA oligos with appropriate 4 base-pair overhangs (IDT) were phosphorylated by incubating with 5 units of T4 PNK for 30 min at 37°C, followed by heating to 95°C and cooling to 25°C at a rate of 0.1°C/s to anneal into duplexes.

Golden Gate reaction mixtures were assembled containing 10 fmol annealed duplex, 25 ng lentiviral backbone, 500 units T7 ligase and 10 units BsmBI-v2. The reaction mix was incubated at 37°C for 5 minutes followed by 5 minutes at 25°C for a total of 15 cycles. The Golden Gate reaction mix was cleaned using DNA Clean and Concentrator columns (Zymo) and transformed into Stbl3 bacteria. Successful cloning was confirmed by Sanger sequencing the resulting plasmids with the hU6-F sequencing primer (see Supplementary Table S7 for sgRNA and primers).

### Large-scale lentivirus production and titer quantification

Lenti-X cells were seeded at 15e6 cells per 150 mm tissue culture dish in antibiotic free complete DMEM media (10% FBS, 1mM sodium pyruvate). After over-night incubation, the cells were treated with 25 uM chloroquine diphosphate for 5 hours. Lentivirus packaging plasmid psPAX2, virus envelope plasmid pCMV-VSV-G and appropriate transfer plasmid were mixed at a molar ratio of 1:0.54:1.26 in 3ml Opti-MEM (Gibco), using a total of 10 pmol plasmid per 150mm plate. Separately, linear polyethylenimine (PEI, 25,000 Da) was mixed with 3 ml OptiMEM at 3 μg PEI per μg total plasmid DNA. The plasmid mixture was gently mixed with the PEI solution and allowed to incubate for 20 min at room temperature, followed by gentle addition to the LentiX cells (6 ml per plate). Following over-night transfection, the media was replaced with virus collection media (10% FBS, 1x penicillin/streptomycin, 1 mM sodium pyruvate in DMEM), and allowed to incubate for 48 hours. Lentivirus supernatant was collected and clarified through centrifugation for 7 minutes at 500xg followed by filtration through a 0.45 μm PES vacuum filter to remove cellular debris. For lentiviral particle concentration, the supernatant was mixed with 8.5% (w/v) polyethylene glycol (PEG, 6,000 Da) and 0.4 M sodium chloride, followed by incubation for 2 hours at 4°C with intermittent mixing. Lentiviral particles were precipitated through centrifugation at 1,500 xg for 45 min at 4°C, supernatant was carefully removed by decanting and the lentivirus-containing pellets were resuspended in sterile PBS at a nominal concentration factor of 100x.

To allow for lentivirus titer quantification, a color-switching Cre-reporter cell line was generated by transducing LentiX cells with CMV-LoxP-DsRed-LoxP-eGFP lentivirus at low MOI and purification of DsRed expressing cells using FACS. This reporter cell line, referred to as LRLG, switches from DsRed fluorescence to eGFP following Cre recombination. To determine Cre lentivirus titer, 500,000 LRLG cells were seed per 12 well in 1ml lentivirus-containing media, supplemented with 8 μg/ml polybrene. Following, 48-hour incubation the fraction of eGFP positive cells in each well was determined using flow cytometry on an Accuri C6 flow cytometer, and original virus titer was back-calculated using the dilution factor and number of seeded cells. To avoid under-, or over-estimation of virus titer, only wells with an eGFP positive fraction of 10-30% were used for calculations. All reagents are listed in Supplementary Table S11.

### Pancreatic organoid establishment and culture

Normal mouse pancreas organoids were established, maintained and propagated as previously described (Boj 2015, Huch 2013). Briefly, a roughly 5×5 mm piece of tumor was collected in sterile HBSS, and finely minced with sterile scalpels. After a 10ml HBSS wash, pancreas pieces were incubated with 8 ml pancreas digestion media (0.4 mg/ml Collagenase P in HBSS) for 20-30 min in a rotating oven at 37°C, until small acinar clusters were observed. The cell pellet was washed once in wash media (DMEM supplemented with 1%FBS and 1x penicillin/streptomycin), followed by washing in serum-free DMEM and seeding in 50ul Matrigel 24-well domes. Following incubation at 37°C for 20 min, Matrigel domes were covered with 0.5 ml complete organoid media supplemented with 10.5 μM ROCK inhibitor (Y-27632). ROCK inhibitor was added only when organoids were seeded or passaged. 0.25 μg/ml Amphotericin B was added in the initial passage to prevent fungal contamination. Complete mouse pancreas organoid media consisted of splitting media (Advanced DMEM/F-12 with 1x GlutaMAX, 10mM HEPES and 1x primocin) supplemented with 0.5 μM TGF-beta inhibitor A83-01, 0.05 μg/ml mEGF, 0.1 μg/ml hFGF-10, 0.1 μg/ml mNoggin, 0.01 μM gastrin I, 1.25 mM N-acetylcysteine, 10 mM nicotinamide, 1x B27 supplement, 50% v/v Wnt3a-conditioned media, and 10% v/v R-spondin I conditioned media. For flow cytometry, transplantation and transduction experiments organoid-containing matrigel domes were disrupted in splitting media containing 2 mg/ml dispase in a rotating oven for 20 min at 37°C. Organoids were then washed once in serum free media and were incubated with TrypLE for approximately 10 minutes with intermittent mechanical disruption. DNaseI was added if clumping was observed. Further digestion was stopped by washing once in wash media, followed by resuspending the cell pellet FACS buffer (2% FBS, 2mM EDTA in PBS) for flow cytometry or serum free DMEM for transplantation. mCherry^+^ expressing cells were sorted on the Aria Cell Sorter II (BD Bioscience). DAPI was used to eliminate dead cells. All reagents are listed in Supplementary Table S11.

### Organoid transduction

Single cell suspensions were mixed with lentiviral particles and polybrene (8 μg/ml) in ultra-low attachment plates. Following over-night incubation, supernatant was replaced with complete organoid media supplemented with 2% Matrigel. After 4 days of culture, puromycin or hygromycin, and 4-hydroxytamoxifen (4OHT) were added. After 10 days of selection, organoid domes were seeded as normal.

### RT-qPCR

Organoid RNA was isolated using Trizol and the miRNeasy kit following the manufacturer’s instructions, using 0.5 ml Trizol per 50 μl Matrigel dome and including the optional on-column DNase treatment step. cDNA was generated using the SuperScript III First-Strand Synthesis System (Invitrogen) and qPCR was performed using Power SYBR Green PCR Master Mix and a QuantStudio3 thermocycler. Primer sequences are listed in Supplementary Table S7. Transcript abundance was calculated as the difference relative to the housekeeping gene bactin (2^−dCT^) and to the control sample (ddCT method). QuantStudio and Excel software was used to analyze the data.

### Immunofluorescence

Immunofluorescence of cultured cells: Cultured cells were fixed in 10% formalin, blocked with 10% normal donkey serum/1% BSA/0.1% saponin, stained for dCas9VP64, MYC, YAP1 and/or γH2AX, followed by conjugated secondary antibodies and DAPI.

Tissue section immunofluorescence: Pancreatic and lung tumor tissues were fixed in zinc/formalin, embedded in paraffin and microtome sectioned. Sections were subjected to citrate antigen retrieval for 15 min at 95C, blocked with 10% normal donkey serum/2% BSA/0.3% Triton X-100, stained for dCas9VP64, MYC and/or YAP1 followed by conjugated secondary antibodies and DAPI (Supplementary Table S12). Autofluorescence was quenched using the TrueVIEW Autofluorescence Quenching Kit and slides were mounted with ProLong Diamond.

Organoid wholemount immunofluorescence: Organoids growing in Matrigel domes were fixed in ice cold 1:1 DMSO:methanol, blocked with 10% normal donkey serum/2% BSA/0.1% saponin, stained for dCas9VP64, Myc, YAP1 followed by conjugated secondary antibodies and DAPI. All reagents are listed in Supplementary Table S11.

### Epifluorescent and confocal imaging

Epifluorescent imaging was performed on an Olympus IX73 inverted fluorescent microscope. Confocal imaging was performed using an Olympus FV1000 laser scanning microscope.

### Quantification of γH2AX foci

Immunofluorescent labeled cells cultured in chamber slides were scanned using a Cytation 5 Imaging Multi-Mode Reader (BioTek). Cell nuclei were identified using DAPI and the abundance of γH2AX foci was determined using CellProfiler (v.4.1.3).

### MTT drug response assay

Cellular drug response was determined as previously described (Thege 2019), briefly, 5000 cells were seeded per 96-well and cells were treated with varying concentrations of oxaliplatin for 96 hours. MTT was added (0.5 mg/ml) for 4 hours, supernatant removed, and formazan crystals dissolved in acidified isopropanol (50 mM HCl, 0.1% Triton X-100). Formazan absorbance was read at 560 nm with 690 nm as reference, on a Cytation 3 Imaging Multi-Mode Reader (BioTek). Oxaliplatin IC50 values were calculated using curve-fitting in Prism (v.9.0.0). All reagents are listed in Supplementary Table S11.

#### Immunoblotting analysis

Primary lung tumor cells, and organoid cell lines were processed for immunoblotting according to a previously described protocol (30). Antibodies are listed in Supplementary Table S12. Blots were analyzed using a ChemiDoc XRS (Bio-Rad). For pharmacological MYC suppression, cells were seeded in 6-well plates and incubated with varying concentrations of MYCi361 for 48 hours followed by immunoblot analysis for MYC and beta-actin. All reagents are listed in Supplementary Table S11.

#### Immunohistochemistry and histology

Immunohistochemistry and H&E staining were performed as previously described. Tissue sections (3–5 μm) sections were stained with hematoxylin and eosin (HE) or used for immunohistochemistry, using standard avidin-biotin histologic methods. Antibodies are listed in Supplementary Table 11. Quantification was done using Aperio ImageScope (v.12.4.3.5008). All reagents are listed in Supplementary Table S11.

### Intranasal instillation and orthotopic pancreatic organoid transplantations

All procedures were performed in a vaporizer applying isoflurane 2% and O2 flow 2L/min. For nasal installation mice taken in and out to administer the virus. All mice were between 8 and 12 weeks of age. Mice of this age are old enough to recover from the anesthetic and the volume of virus administered to the lung. Intranasal instillations have been described previously(17). In brief, mice were transduced with a total volume of 80 μl per mouse. After mice were anaesthetized (to retain deep inhalation), mice were placed on their back and the virus solution was administered dropwise and slowly into one nostril until virus was completely inhaled. Breathing of mice was monitored until fully awake. Lung tissues were collected, fixed, embedded in paraffin and stained with H&E, for YFP, MYC, YAP1, dCas9VP64, and different immune cell, proliferation, and apoptosis marker as described above. For dose dependent transfection efficiency and to confirm long-term stable transfection GFP/YFP/Cre positive cells in pancreatic tissues 10 and 28 days after virus delivery were quantified using APERIO/image scope. For survival time point, tumor incidence and tumor size were quantified and representative macroscopic and microscopic H&E and YFP/mCherry pictures were taken. Orthotopic transplantations of pancreatic organoids were performed as previously described (Boj 2015). In brief, 5×10^5^ organoid cells prepared as single cell solutions as described above were injected into the pancreas of C57BL/6NJ PPKS syngeneic littermates using a 26 G needle. The mice were sacrificed at survival timepoint, defined when the mice reached sickness criteria. Tumor incidence and tumor sizes were quantified in a blinded manner and representative macroscopic (and mCherry+) and microscopic images were taken.

### Tumor mRNA-seq and expression profiling

Total RNA was isolated from 16 mouse lung tumors (4 PPKY/LV, 4 PPKS/NT, 4 PPKS/M and 4 PPKS/Y) using the AllPrep DNA/RNA Mini Kit (Qiagen). Indexed libraries were generates using the KAPA mRNA HyperPrep kit and sequenced on a NextSeq500 sequencer (NextSeq 500/550 High Output Kit v2.5, 75 SE). Adapter trimming and FASTQ generation was done with Basespace (Illumina). Reads were aligned to mouse reference genome (mm10) and differential expression was determined using the RNA-Seq Alignment (v.2.0.2) and RNA-Seq Differential Expression (v1.0.1) Basespace applications (Illumina). Gene Set Enrichment Analysis (GSEA) was performed using the WebGestalt online portal (http://www.webgestalt.org/) and GSEAJava (v.4.1.0). Data visualization was performed in R (v.3.6.0) using Rstudio (v.1.1.463). Heatmaps were generated using heatmap.2 (gplots, v.3.6.3). GSEA dot plots were generated with ggplot2 (v.3.3.3). Raw data were deposited in ArrayExpress, accession E-MTAB-11122.

### Mouse tumor transcriptomic subtyping

Mouse lung tumors were subtyped using CMScaller (v.0.9.2, Eide 2017). Lists of subtype-specific genes were downloaded from Jang et al. Orthologs for over-expressed genes from each subtype were generated using DIOPT (v.8.5, https://www.flyrnai.org/cgi-bin/DRSC_orthologs.pl), and can be found in Supplementary Table 6. Only up-regulated genes from each subtype were used for CMScaller subtyping. Gene symbol, entrez id and expression value (rlog) for each sample was used as CMScaller input.

### TCGA bioinformatics

Gene expression and copy number data from the TCGA-LUAD study were downloaded from Xena (http://xena.ucsc.edu/). TCGA-LUAD tumors were subtyped using CMScaller (v.0.9.2, Eide 2017). Lists of subtype-specific over-expressed genes were downloaded from Jang et al. and can be found in Supplementary Table 6. Only up-regulated genes from each subtype were used for CMScaller subtyping. Gene symbol, entrez id and expression value (fpkm-uq) for each sample was used as CMScaller input. Differential Expression was performed in R using DESeq2 (v.1.26.0). GSEA for MYC-associated gene sets was performed using GSEAJava (v.4.1.0). Data visualization was performed in R (v.3.6.0) using Rstudio (v.1.1.463). Heatmaps were generated using heatmap.2 (gplots, v.3.6.3).

## Schematics

Schematics were generated using BioRender (https://biorender.com/).

## Data availability

Raw lung tumor mRNA-seq data were deposited in ArrayExpress, accession E-MTAB-11122.

## Code availability

The codes supporting the current study are available from the corresponding author on request.

## Materials availability

Plasmids generated in this study are available on request.

## QUANTIFICATION AND STATISTICAL ANALYSIS

Description of sample size (n) and statistical tests are detailed in the figure legends or methods. All values for n are for individual mice or individual sample (biological replicates) unless otherwise noted. Sample sizes were chosen based on previous experience with given experiments. Data are expressed as mean ± SEM unless otherwise noted. Differences were analyzed by two-sided Student’s t-test, or by one-way ANOVA with Tukey’s multiple comparison test. The Log-rank (Mantel-Cox) test was applied to determine survival significance and the Pearson r and two-sided test calculated for correlations. Statistical analysis was performed with Prism (GraphPad, v.9.0.0). . p-values ≤ 0.05 were considered significant.

## Acknowledgements

We thank members of the Rhim and Maitra laboratories for critical reading of the manuscript.

F.I.T. is supported by the Ben and Rose Cole Charitable Pria Foundation, S.M.W. is supported by German Cancer Aid, the Mildred-Scheel-Postdoctoral Fellow program, the Ben and Rose Cole Charitable Pria Foundation, the MD Anderson GI SPORE career enhancement program. A.D.R. is supported by the V Foundation (V Scholar Grant), Doris Duke Foundation (Clinical Scholar Grant), Andrew Sabin Family Foundation, Cockrell Family Foundation, NCI (MD Anderson SPORE), MD Anderson Cancer Center (Physician Scientist Program, Pancreatic Cancer Moonshot) and CPRIT (Rising Stars Award). A.M. is supported by the Khalifa bin Zayed Foundation and the MD Anderson Moonshot Program in Pancreatic Cancer. The MDACC Flow cytometry and cellular imaging core facility is partially funded by the NCI Cancer Center Support Grant P30CA16672. The MDACC Genetically Engineered Mouse Facility (GEMF) is partially funded by NCI Cancer Center Support Grant P30CA016672 and NCI R50CA211121.

## Author contributions

F.I.T. conceived of the project, F.I.T. and S.M.W. designed and performed experiments, and analyzed the data, D.N.R., B.B.B. and S.L.M. performed experiments, A.M. revised and edited the manuscript, A.R. edited the manuscript, F.I.T. and S.M.W. wrote and revised the manuscript.

---

**Supplementary Figure 1 (related to Figure 1).**
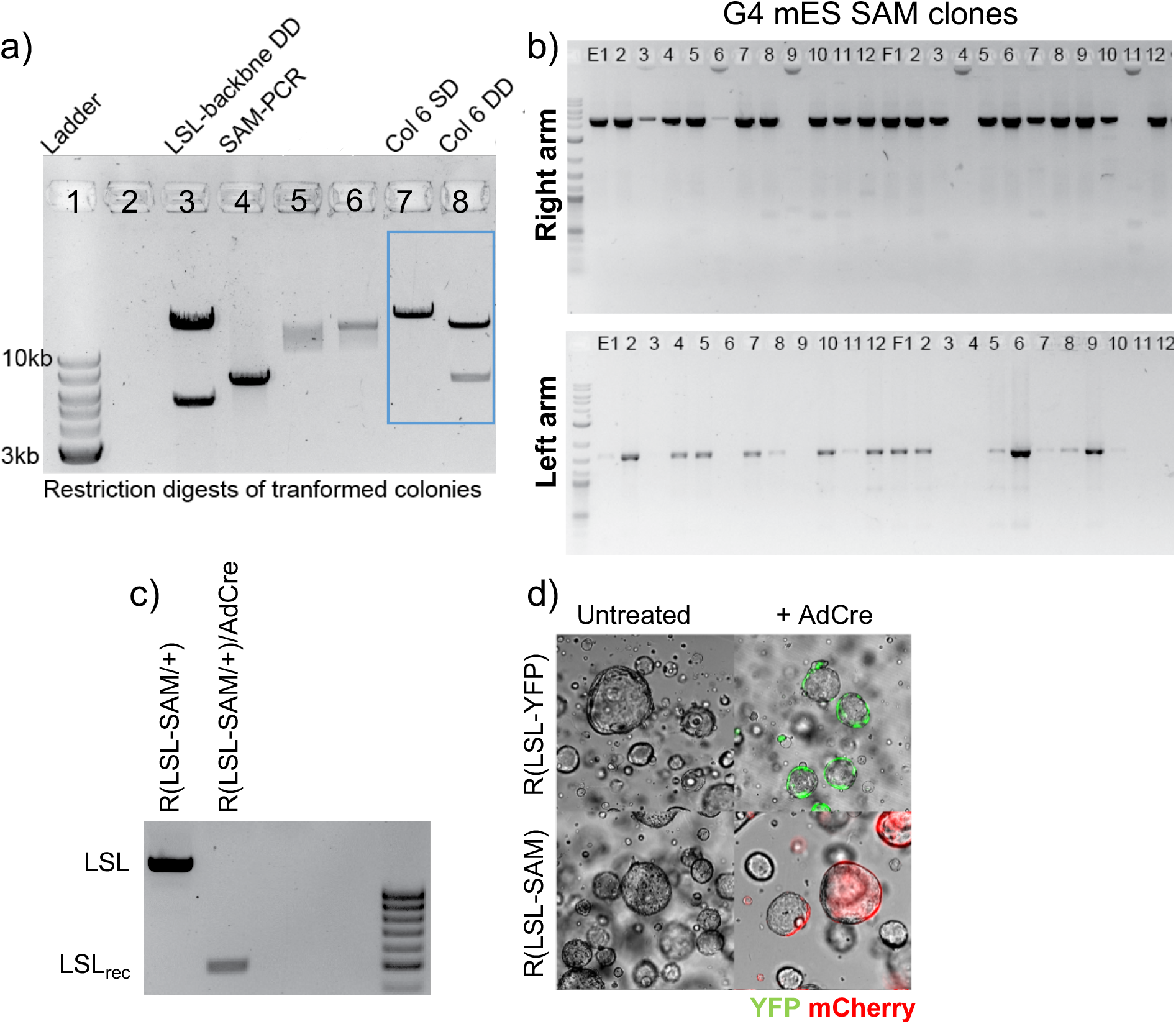
Development and validation of the LSL-SAM mouse model. **(a)** Restriction digestion analysis of cloned CAG-LSL-dCas9VP64-P2A-MS2-p65-HSF1-T2A-mCherry-Rosa26TV constructs. Lane 3 and 4 containing starting material and lane 7 and 8 containing correctly assembled clone. (SD– single digest, DD – double digest) **(b)** PCR confirmation of LSL-SAM Rosa26 knock-in (right and left homology arm) in G4 mouse embryonic stem cells. **(c)** PCR reaction confirming LSL cassette recombination in FACS sorted R26(LSL-SAM/+) pancreatic organoids transduced with adenovirus Cre (AdCre). **(d)** Fluorescent imaging of AdCre transduced and untreated pancreatic organoids from R26(LSL-SAM/+) and R26(LSL-YFP/+) mice.

**Supplementary Figure 2 (related to Figure 2).**
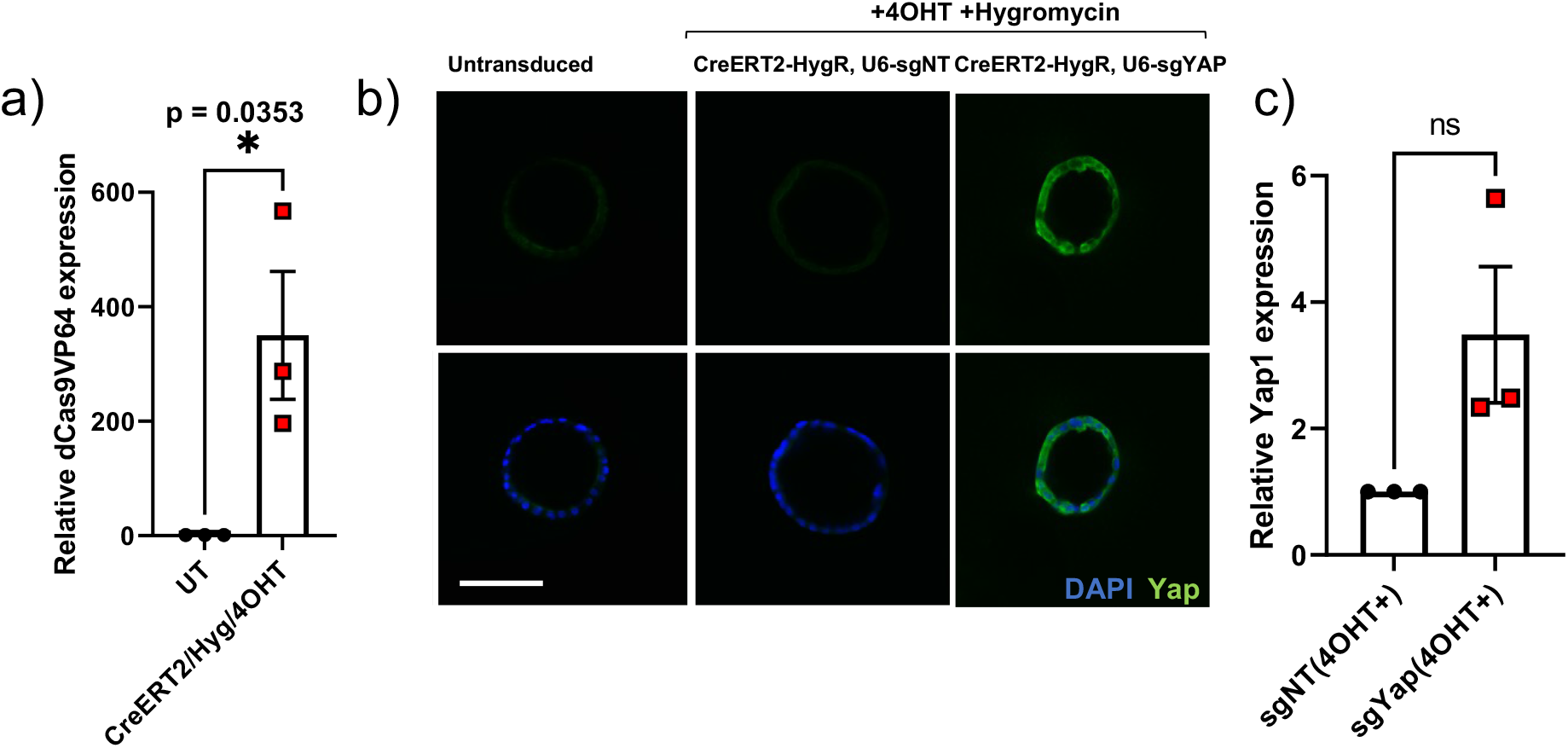
Functional validation of the LSL-SAM/CRISPRa construct in pancreas organoids. **(a)** qPCR analysis showing expression of dCas9VP64 in P53(F/F), Kras(LSL-G12D), Rosa26(LSL-SAM); PPKS pancreas organoids following transduction with CMV-CreERT2-P2A-hygR lentivirus, selection and treatment with 4OHT, relative to untreated control, unpaired, two-tailed, Student’s t-test. **(b)** Wholemount immunofluorescent staining for YAP1 in PPKS pancreatic organoids transduced with *Yap1*-targeted or non-targeted CMV-CreERT2-P2A-hygR-U6-sgRNA(MS2) lentivirus and untransduced controls, YAP1 (green), DAPI (blue), scale bar 100 μm. **(c)** qPCR analysis showing *Yap1* gene activation in PPKS pancreatic organoids transduced with *Yap1*-targeted CMV-CreERT2-P2A-hygR-U6-sgRNA(MS2) lentivirus relative to non-targeted control, unpaired, two-tailed, Student’s t-test.

**Supplementary Figure 3 (related to Figure 3).**
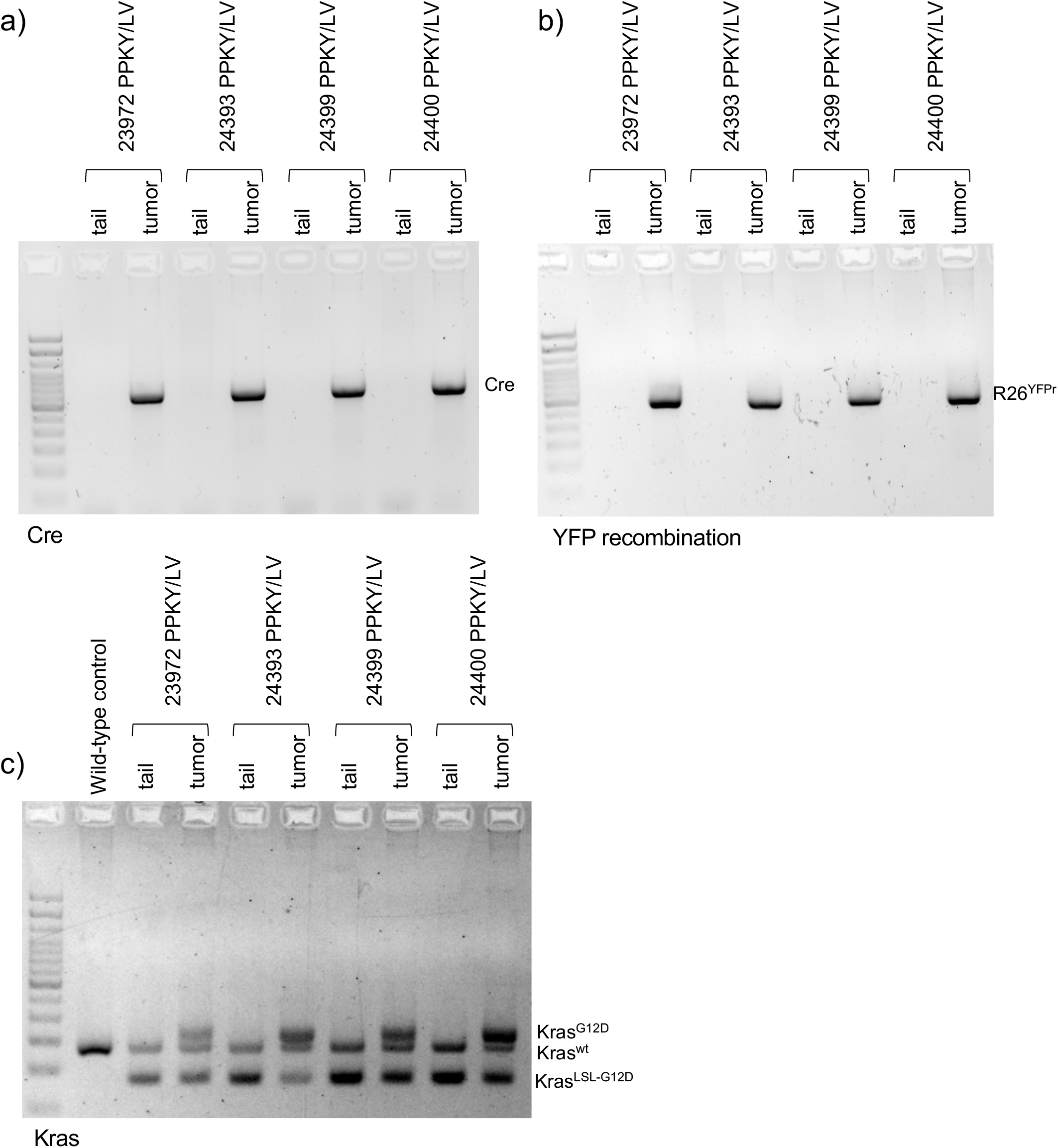
Retained viral payload and Cre-mediated recombination in transduction initiated lung tumors. **(a)** PCR analysis of viral payload (Cre sequence) in lung tumor DNA from CMV-Cre transduction-initiated tumors in P53(F/F), Kras(LSL-KrasG12D/+), Rosa26(LSL-YFP); PPKY mice, with somatic (tail) DNA from the same mouse as control. **(b)** YFP recombination in lung tumor DNA from CMV-Cre transduction-initiated tumors in P53(F/F), Kras(LSL-KrasG12D/+), Rosa26(LSL-YFP); PPKY mice, with somatic (tail) DNA from the same mouse as control. YFPr – recombined LSL-YFP cassette. **(c)** Kras recombination in lung tumor DNA from CMV-Cre transduction-initiated tumors in P53(F/F), Kras(LSL-KrasG12D/+), Rosa26(LSL-YFP); PPKY mice, with somatic (tail) DNA from the same mouse as control. (wt) – wild type allele, (LSL-G12D) – unrecombined oncogenic Kras cassette, (G12D) – recombined oncogenic Kras cassette.

**Supplementary Figure 4 (related to Figure 4).**
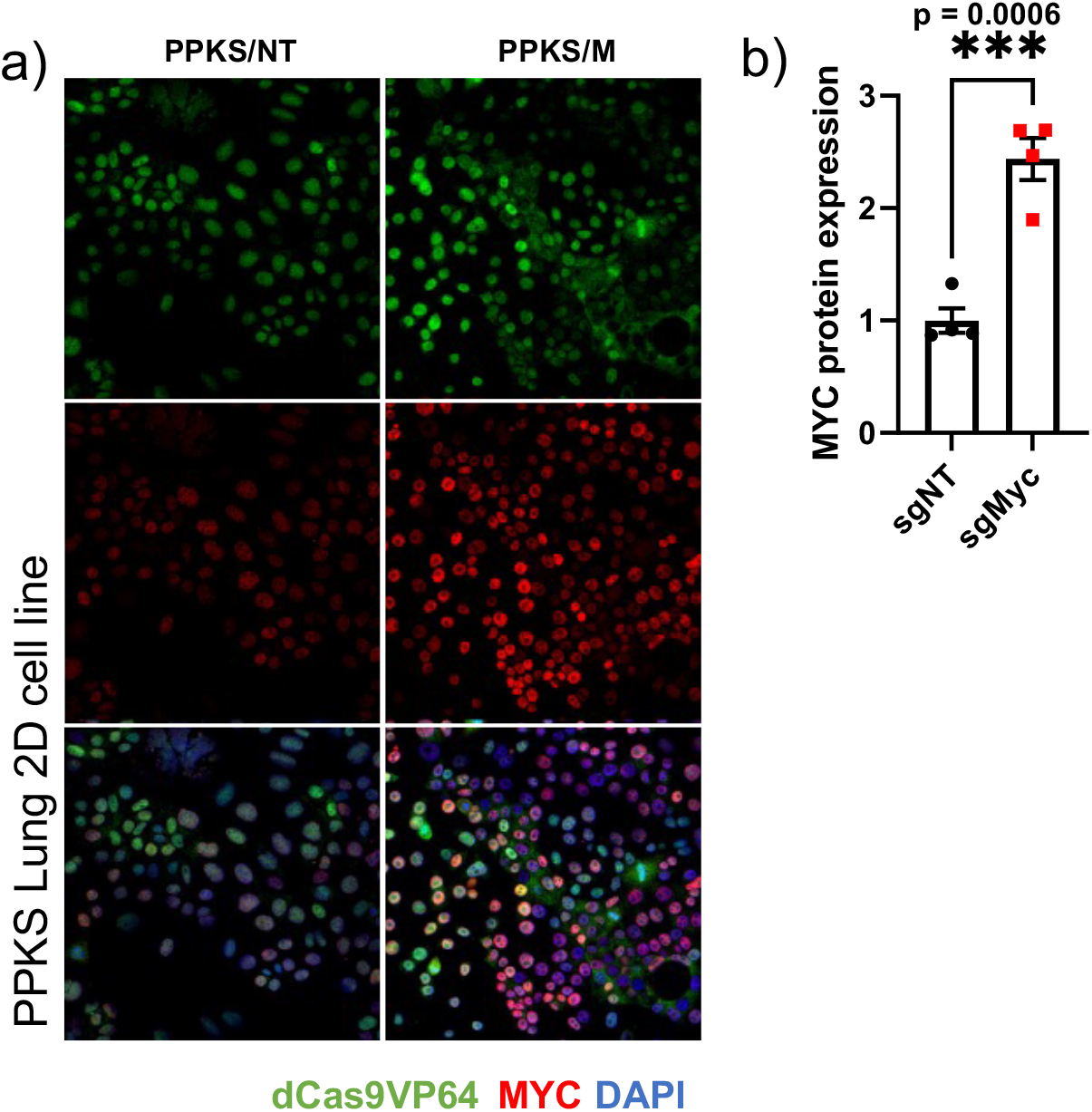
MYC over-expression in cell lines derived from *Myc*-activated lung tumors (PPKS/M). **(a)** Immunofluorescent staining of dCas9VP64 and MYC in cell lines derived from *Myc*-activated (PPKS/M) and non-targeted control (PPKS/NT) lung tumors, revealing nuclear MYC-overexpression in PPKS/M tumor-derived cell lines relative to controls and retained dCas9VP64 expression in both cell lines. dCas9VP64 (green), MYC (red), DAPI (blue). **(b)** Quantification of immunoblot analysis of dCas9VP64, MYC and ß-actin in cell lines derived from *Myc*-activated (PPKS/M) and non-targeted control (PPKS/NT) lung tumors, revealing 2.5-fold average MYC-overexpression in PPKS/M tumor-derived cell lines relative to controls

**Supplementary Figure 5 (related to Figure 6).**
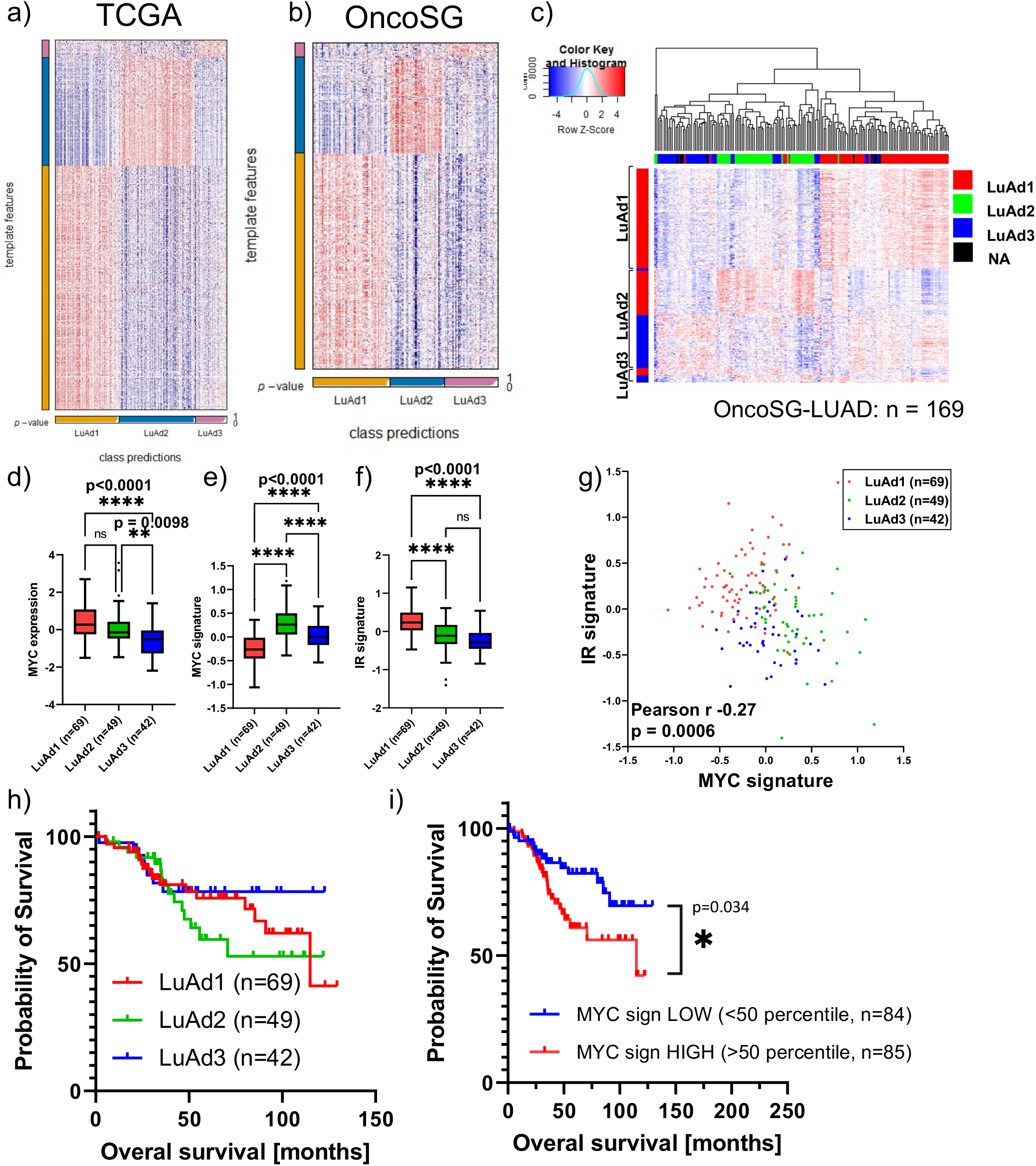
Transcriptomic subtyping of TCGA-LUAD and OncoSG lung adenocarcinoma tumors. **(a)** CMScaller classification heatmap of the TCGA-LUAD data set. **(b)** CMScaller classification heatmap of the OncoSG data set. **(c)** Heatmap and non-supervised hierarchical clustering of OncoSG patient tumors based on subtype-specific gene expression. **(d)** Normalized *MYC* expression in LuAd1, LuAd2 and LuAd3 tumors from the OncoSG data set, one-way ANOVA with Tukey’s multiple comparison test. **(e)** MYC signature expression in LuAd1, LuAd2 and LuAd3 tumors from the OncoSG data set, one-way ANOVA with Tukey’s multiple comparison test. **(f)** IR-signature as function of LuAd subtype in the OncoSG data set, one-way ANOVA with Tukey’s multiple comparison test. **(g)** Correlation plot of IR signature and MYC signature in patient tumors from the OncoSG data set, Pearson’s correlation coefficient (r = −0.26, p=0.0006). **(h)** Kaplan-Meier survival plot of the OncoSG data set, stratified by patient tumor subtype; LuAd1 (n=69), LuAd2 (n=49) and LuAd3 (n=42). **(i)** Kaplan-Meier survival plot of the OncoSG data set, stratified by patient tumor MYC-signature; MYC-signature LOW (<50 percentile, n=84), HIGH (>50 percentile, n=85).

## References

1. Jinek M, Chylinski K, Fonfara I, Hauer M, Doudna JA, Charpentier E. A programmable dual-RNA-guided DNA endonuclease in adaptive bacterial immunity. Science 2012;337:816–21

2. Cong L, Ran FA, Cox D, Lin S, Barretto R, Habib N, et al. Multiplex genome engineering using CRISPR/Cas systems. Science 2013;339:819–23

3. Mali P, Yang L, Esvelt KM, Aach J, Guell M, DiCarlo JE, et al. RNA-guided human genome engineering via Cas9. Science 2013;339:823–6

4. Platt RJ, Chen S, Zhou Y, Yim MJ, Swiech L, Kempton HR, et al. CRISPR-Cas9 knockin mice for genome editing and cancer modeling. Cell 2014;159:440–55

5. Chiou SH, Winters IP, Wang J, Naranjo S, Dudgeon C, Tamburini FB, et al. Pancreatic cancer modeling using retrograde viral vector delivery and in vivo CRISPR/Cas9-mediated somatic genome editing. Genes Dev 2015;29:1576–85

6. Maresch R, Mueller S, Veltkamp C, Ollinger R, Friedrich M, Heid I, et al. Multiplexed pancreatic genome engineering and cancer induction by transfection-based CRISPR/Cas9 delivery in mice. Nat Commun 2016;7:10770

7. Mueller S, Engleitner T, Maresch R, Zukowska M, Lange S, Kaltenbacher T, et al. Evolutionary routes and KRAS dosage define pancreatic cancer phenotypes. Nature 2018;554:62–8

8. Konermann S, Brigham MD, Trevino AE, Joung J, Abudayyeh OO, Barcena C, et al. Genome-scale transcriptional activation by an engineered CRISPR-Cas9 complex. Nature 2015;517:583–8

9. Chavez A, Tuttle M, Pruitt BW, Ewen-Campen B, Chari R, Ter-Ovanesyan D, et al. Comparison of Cas9 activators in multiple species. Nat Methods 2016;13:563–7

10. Joung J, Konermann S, Gootenberg JS, Abudayyeh OO, Platt RJ, Brigham MD, et al. Genome-scale CRISPR-Cas9 knockout and transcriptional activation screening. Nat Protoc 2017;12:828–63

11. Fidanza A, Lopez-Yrigoyen M, Romano N, Jones R, Taylor AH, Forrester LM. An all-in-one UniSam vector system for efficient gene activation. Sci Rep 2017;7:6394

12. Hunt C, Hartford SA, White D, Pefanis E, Hanna T, Herman C, et al. Tissue-specific activation of gene expression by the Synergistic Activation Mediator (SAM) CRISPRa system in mice. Nat Commun 2021;12:2770

13. Naldini L, Blomer U, Gallay P, Ory D, Mulligan R, Gage FH, et al. In vivo gene delivery and stable transduction of nondividing cells by a lentiviral vector. Science 1996;272:263–7

14. Kumar MS, Erkeland SJ, Pester RE, Chen CY, Ebert MS, Sharp PA, et al. Suppression of non-small cell lung tumor development by the let-7 microRNA family. Proc Natl Acad Sci U S A 2008;105:3903–8

15. Jackson EL, Willis N, Mercer K, Bronson RT, Crowley D, Montoya R, et al. Analysis of lung tumor initiation and progression using conditional expression of oncogenic K-ras. Genes Dev 2001;15:3243–8

16. Meuwissen R, Linn SC, van der Valk M, Mooi WJ, Berns A. Mouse model for lung tumorigenesis through Cre/lox controlled sporadic activation of the K-Ras oncogene. Oncogene 2001;20:6551–8

17. DuPage M, Dooley AL, Jacks T. Conditional mouse lung cancer models using adenoviral or lentiviral delivery of Cre recombinase. Nat Protoc 2009;4:1064–72

18. Kortlever RM, Sodir NM, Wilson CH, Burkhart DL, Pellegrinet L, Brown Swigart L, et al. Myc Cooperates with Ras by Programming Inflammation and Immune Suppression. Cell 2017;171:1301–15 e14

19. Topper MJ, Vaz M, Chiappinelli KB, DeStefano Shields CE, Niknafs N, Yen RC, et al. Epigenetic Therapy Ties MYC Depletion to Reversing Immune Evasion and Treating Lung Cancer. Cell 2017;171:1284–300 e21

20. Casey SC, Tong L, Li Y, Do R, Walz S, Fitzgerald KN, et al. MYC regulates the antitumor immune response through CD47 and PD-L1. Science 2016;352:227–31

21. Han H, Jain AD, Truica MI, Izquierdo-Ferrer J, Anker JF, Lysy B, et al. Small-Molecule MYC Inhibitors Suppress Tumor Growth and Enhance Immunotherapy. Cancer Cell 2019;36:483–97 e15

22. Jang HJ, Lee HS, Ramos D, Park IK, Kang CH, Burt BM, et al. Transcriptome-based molecular subtyping of non-small cell lung cancer may predict response to immune checkpoint inhibitors. J Thorac Cardiovasc Surg 2020;159:1598–610 e3

23. Schaub FX, Dhankani V, Berger AC, Trivedi M, Richardson AB, Shaw R, et al. Pan-cancer Alterations of the MYC Oncogene and Its Proximal Network across the Cancer Genome Atlas. Cell Syst 2018;6:282–300 e2

24. Kidder BL, Yang J, Palmer S. Stat3 and c-Myc genome-wide promoter occupancy in embryonic stem cells. PLoS One 2008;3:e3932

25. Kim J, Chu J, Shen X, Wang J, Orkin SH. An extended transcriptional network for pluripotency of embryonic stem cells. Cell 2008;132:1049–61

26. Chandriani S, Frengen E, Cowling VH, Pendergrass SA, Perou CM, Whitfield ML, et al. A core MYC gene expression signature is prominent in basal-like breast cancer but only partially overlaps the core serum response. PLoS One 2009;4:e6693

27. Masso-Valles D, Beaulieu ME, Soucek L. MYC, MYCL, and MYCN as therapeutic targets in lung cancer. Expert Opin Ther Targets 2020;24:101–14

28. Eide PW, Bruun J, Lothe RA, Sveen A. CMScaller: an R package for consensus molecular subtyping of colorectal cancer pre-clinical models. Sci Rep 2017;7:16618

29. Rhim AD, Mirek ET, Aiello NM, Maitra A, Bailey JM, McAllister F, et al. EMT and dissemination precede pancreatic tumor formation. Cell 2012;148:349–61

30. Ai J, Wormann SM, Gorgulu K, Vallespinos M, Zagorac S, Alcala S, et al. Bcl3 Couples Cancer Stem Cell Enrichment With Pancreatic Cancer Molecular Subtypes. Gastroenterology 2021

